# Syngap1 Dynamically Regulates the Fine-Scale Reorganization of Cortical Circuits in Response to Sensory Experience

**DOI:** 10.1101/2020.04.14.041038

**Authors:** Nerea Llamosas, Thomas Vaissiere, Camilo Rojas, Sheldon Michaelson, Courtney A. Miller, Gavin Rumbaugh

**Author notes:** Corresponding author: Gavin Rumbaugh 130 Scripps Way, #3B3 Jupiter, FL, 33458.

## Abstract

Experience induces complex, neuron-specific changes in population activity within sensory cortex circuits. However, the mechanisms that enable neuron-specific changes within cortical populations remain unclear. To explore the idea that synapse strengthening is involved, we studied fine-scale cortical plasticity in *Syngap1* mice, a neurodevelopmental disorder model useful for linking synapse biology to circuit functions. Repeated functional imaging of the same L2/3 somatosensory cortex neurons during single whisker experience revealed that *Syngap1* selectively regulated the plasticity of a low-active, or “silent”, neuronal subpopulation. *Syngap1* also regulated spike-timing-dependent synaptic potentiation and experience-mediated *in vivo* synapse bouton formation, but not synaptic depression or bouton elimination in L2/3. Adult re-expression of *Syngap1* restored plasticity of “silent” neurons, demonstrating that this gene controls dynamic cellular processes required for population-specific changes to cortical circuits during experience. These findings suggest that abnormal experience-dependent redistribution of cortical population activity may contribute to the etiology of neurodevelopmental disorders.

## Introduction

Sensory experience drives plasticity leading to changes in cortical function (Bavelier et al., 2010; Holtmaat and Svoboda, 2009; Yu and Zuo, 2011). This is a fundamental neural process believed to enhance computational features of cortical networks that process sensory information, which can facilitate behavioral adaptations in response to a changing environment (Greenwood and Parasuraman, 2010; May, 2011; Pascual-Leone et al.; Reed et al., 2011). Experience-dependent plasticity within cortical networks occurs at multiple levels (Feldman and Brecht, 2005). Sensory experience induces plasticity at the level of receptive fields within large brain areas (termed “map plasticity”), leading to macroscopic changes in cortical function (Polley et al., 2004). Sensory map plasticity is mediated by fine-scale changes at the level of individual neurons and the local circuits they comprise. Recent studies suggest that experience influences the function of individual neurons within local sensory cortex networks in complex ways, with spatially intermingled neurons undergoing either up- or down-regulation of activity after sensory deprivation (Margolis et al., 2014). This important finding highlights how fine-scale redistribution of population activity is accompanied by retuning of individual neurons to sensory input, providing a tangible example of how experience transforms the computational properties of cortical networks.

Experience engages diverse cellular mechanisms that act in concert to cause complex changes in cortical population activity. For example, sensory experience leads to the induction of Hebbian-type synaptic plasticity that can strengthen or weaken excitatory synaptic input onto sensory-responsive neurons (Feldman and Brecht, 2005). However, experience-dependent circuit plasticity is not limited to changes in excitatory synaptic strength. Robust changes to the function and connectivity of GABAergic interneurons also occurs in response to novel experience, which in turn regulates the output of pyramidal neurons within cortical microcircuits (Gainey and Feldman, 2017; Griffen et al., 2014; Maffei et al., 2006). Moreover, intrinsic changes to neuronal excitability have also been observed, and in combination with changes to GABAergic function, these processes are thought to maintain overall firing rates within the network even when activity is redistributed among individual neurons in response to sensory experience (Desai et al., 1999; Lambo and Turrigiano, 2013; Margolis et al., 2012). Currently, it remains unclear how these different modes of experience-dependent plasticity contribute to heterogeneous neuron-specific changes within a broader cortical population.

Understanding how cortical connectivity changes during experience is also critical to elucidate the etiology of brain disorders defined by cognitive impairments. Neurodevelopmental disorders (NDDs) with cognitive impairment, such as Autism Spectrum Disorder (ASD), are characterized by robust alterations in functional connectivity (Hahamy et al., 2015; Just et al., 2019). However, similarly robust changes in brain anatomy are less apparent (Haar et al., 2016). These findings suggest that impaired brain function in ASD and related NDDs arise, at least in part, through alterations in how experience sculpts and modifies the fine-scale connectivity of individual neurons within functional cortical networks.

In this study, we sought to understand the mechanisms that underlie fine-scale plasticity required for the redistribution of neuronal population responses during experience. We hypothesized that major NDD risk genes that directly regulate synapse biology enable population-specific plasticity with cortical circuits. To begin to test this idea, we utilized serial *in vivo* two-photon imaging approaches paired with a single whisker experience (SWE) paradigm in *Syngap1* haploinsufficient mice. We chose SWE because it engages multiple cellular mechanisms within cortical circuits to drive complex changes in cortical population responses (Feldman, 2009; Margolis et al., 2012). *Syngap1* haploinsufficient mice are an established model useful for exploring relationships between NDD-mediated genetic risk and forebrain excitatory synapse biology (Clement et al., 2012; Kilinc et al., 2018). Indeed, *de novo* variants leading to *SYNGAP1* haploinsufficiency in humans cause a complex NDD defined by global developmental delay, autistic features, cognitive impairment, and early-onset epilepsy (Holder et al., 2018; Satterstrom et al., 2020; Vlaskamp et al., 2019). This gene directly regulates excitatory synapse structure and function by gating NMDA receptor-dependent regulation of AMPA receptor trafficking and dendritic spine size (Rumbaugh et al., 2006; Vazquez et al., 2004). *Syngap1* heterozygous mice, which are a construct-valid model for the human disorder, have well-documented impairments in dendritic spine synapse structure and function (Aceti et al., 2015; Clement et al., 2012). However, it is not known how *Syngap1*, or any major NDD gene, shapes complex circuit-level plasticity mechanisms required to engage neuron-specific changes in cortical function during sensory experience. Here, we found that *Syngap1* was required for the experience-dependent potentiation of “silent” neurons (Margolis et al., 2012) within L2/3 somatosensory cortex. *Syngap1* was also required for spike-timing-dependent synaptic potentiation and SWE-mediated synapse bouton formation in this same area, but not synaptic depression or synapse bouton elimination. Remarkably, adult re-expression of *Syngap1* restored SWE-mediated potentiation of the L2/3 “silent” neuronal subpopulation. Thus, *Syngap1* contributes to fine-scale sensory map plasticity by promoting experience-dependent increases in synaptic connectivity leading to potentiation of L2/3 population responses. These data suggest that synapse strengthening mechanisms are crucial for redistributing activity within L2/3 “silent” wS1 neurons during experience. They also suggest that genes causally-linked to complex NDDs may impair cortical function by disrupting how experience reorganizes synaptic inputs at the level of individual neurons.

## Results

### Syngap1 is required for population response plasticity within “silent” somatosensory cortex L2/3 neurons

SWE induces cell type-specific changes in cortical population responses. Neurons with reduced sensitivity to whisker stimulation scale-up activity after SWE, while neighboring neurons that are strongly responsive to the same stimulation scale down their activity (Margolis et al., 2012). Thus, SWE is well-suited for understanding how genes linked to synapse function and cognition, such as *Syngap1*, regulate the fine-scale reorganization of cortical circuits and the resulting impacts on neuronal population responses during sensory experience. We first set out to replicate the finding that individual neuronal subpopulations within the same local network differentially respond to SWE (Margolis et al., 2012). Therefore, we initially sought to: *1)* determine if SWE drives subpopulation-specific changes in neuronal responses in *Syngap1^+/+^* (WT) mice, and *2)* understand if and how this form of population-based plasticity is altered by *Syngap1* heterozygosity. Crossing *Syngap1*^+/−^ mice (Clement et al., 2012; Kim et al., 2003) to a line that stably expresses the Thy1-GCaMP6s transgene (GP4.3) (Dana et al., 2014) enabled longitudinal activity measurements of the exact same *w*S1 (whisker somatosensory cortex) L2/3 neuronal population before and after SWE (Fig. 1A). In each animal, we identified two distinct whisker receptive areas of cortex by intrinsic optical signal imaging (IOS), such as β and C2. However, tracking changes to cortical population responses during SWE requires that the neuronal population is stable across multiple baseline sessions (i.e. before trimming). Therefore, we first assessed the stability of baseline neuronal responses evoked during whisker deflections in both genotypes. Responses of L2/3 neurons imaged from both receptive areas were combined and then clustered into three subpopulations based on their response amplitude during baseline imaging sessions (Margolis et al., 2012) (e.g. low, medium, high; Fig. 1B). Importantly, we observed that GCaMP6s responses of individual neurons were stable in each of the three subpopulations during the baseline period in both *Syngap1*^+/+^ (Fig. 1C) and *Syngap1*^+/−^ mice (Fig. 1D).

**Figure 1.**
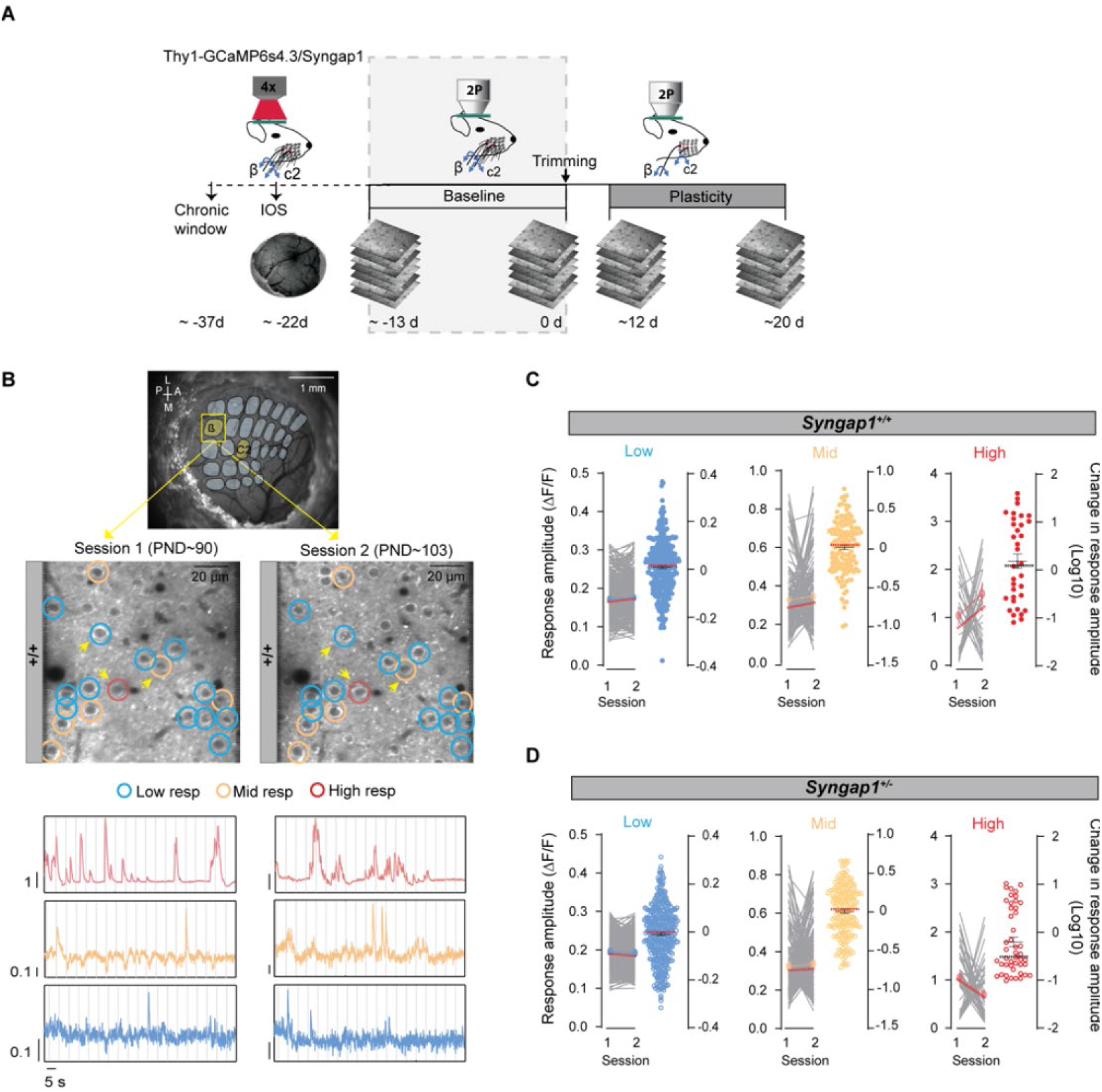
Stable whisker responsiveness of L2/3 neurons under baseline conditions. **(A)** Experimental timeline/design. Thy1-Gcamp6s4.3/*Syngap1*^+/+^ and Thy1-Gcamp6s4.3/*Syngap1*^+/−^ mice were implanted with a chronic cranial window and allowed to recover for at least two weeks before Intrinsic Optical Signal (IOS) imaging. IOS was used to identify the locations activated by the deflection of the principal whisker (PW) and non-principal whisker (NPW), or β and C2 whiskers, respectively. Baseline imaging sessions (grey shaded region) were carried out ~-13 and 0 days before whisker trimming (i.e. day 0). (**B**) Image of cortical surface (top) through a chronic cranial window with an overlaid barred field, imaging areas indicated in yellow (β and C2). Representative in vivo two-photon microscopy images and representative ΔF/F traces (bottom) from the same cells of a *Syngap1^+/+^* mouse over two baseline imaging sessions. Circles show low-(blue), mid- (beige) and high- (red) active cells. (C-D) ΔF/F (± SEM) for each cell class from both β and C2 receptive areas over two baseline sessions in *Syngap1^+/+^* (**C**) and *Syngap1^+/−^* mice (**D**) (n=306 low-active cells, n=143 mid-active cells, n=34 high-active cells from 4 *Syngap1^+/+^* mice; n=414 low-active cells, n=194 mid-active cells, n=46 high-active cells from 6 *Syngap1^+/−^* mice). p > 0.05, Wilcoxon matched-pairs signed rank test. Medians are represented by red or black dashed lines.

The response stability of the three neuronal subpopulations during baseline sessions enabled an analysis of how SWE impacted response magnitude in these same neuronal subpopulations (Fig. 2A). In general, we observed subpopulation-specific changes in neuronal activity during SWE in both genotypes (Fig. 2B, C). Specifically, SWE drove low-active, or “silent”, neurons (Margolis et al., 2012) residing in the spared cortical area (i.e. region of cortex corresponding to the untrimmed whisker) to significantly potentiate their activity in response to non-principal whisker (NPW - trimmed) stimulation in *Syngap1*^+/+^ mice (Fig. 2B, D, E, F). In contrast, spatially intermingled high-active neurons significantly decreased their activity in response to the same whisker stimulation (Fig. 2B, E). In *Syngap1*^+/+^ mice, we also observed scaling-down of the high-active neurons (Fig. 2B, E). However, the low-active “silent” population failed to potentiate in these animals (Fig. 2B, E, F). Because low-active neurons represent the largest subpopulation (i.e. >60% of recorded neurons), the lack of plasticity in this population in *Syngap1*^+/−^ mice drove a large genotype-effect on combined L2/3 wS1 population response dynamics (Fig. 2G).

**Figure 2.**
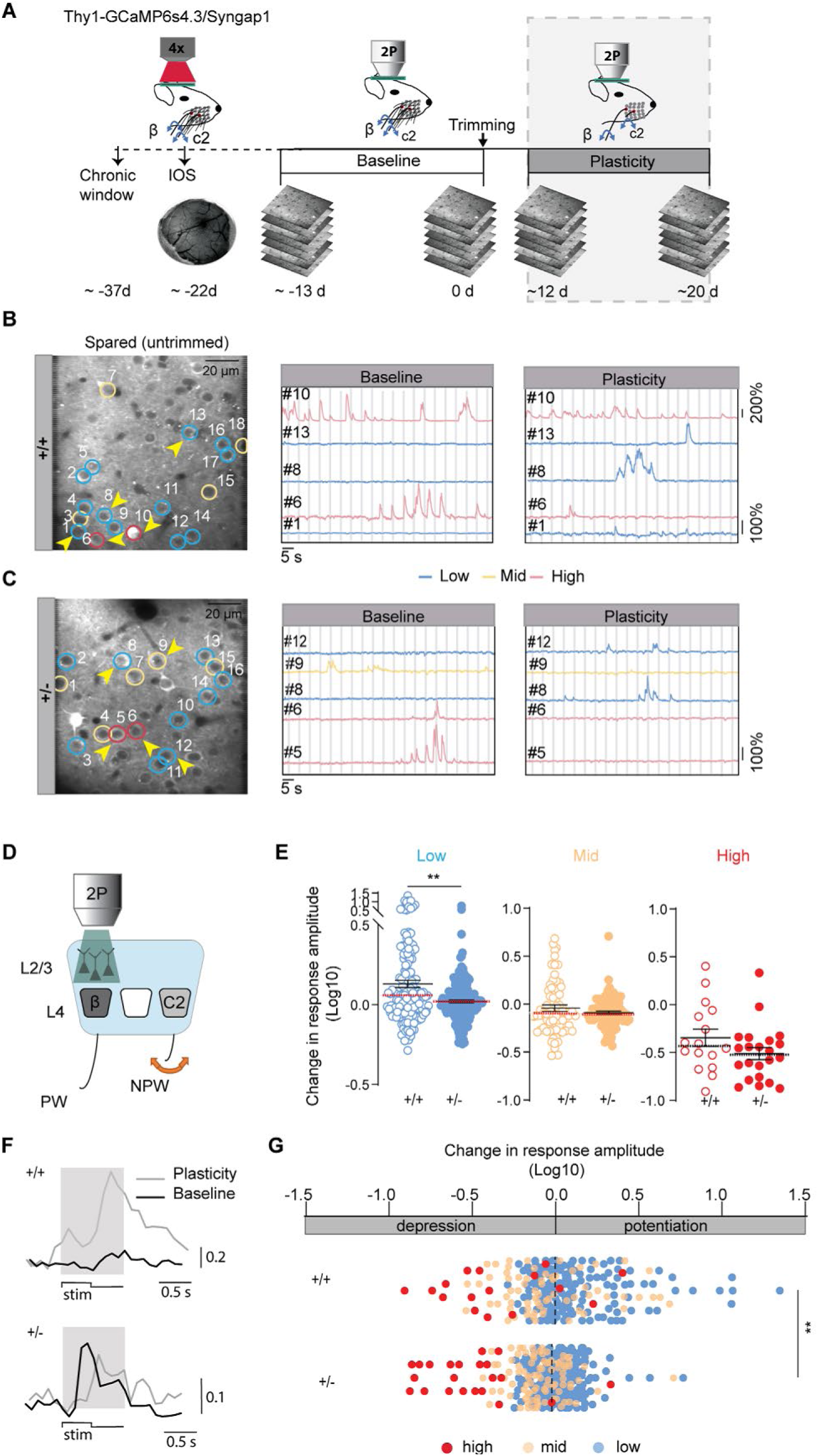
*Syngap1* heterozygosity decreases plasticity of a ‘silent’ neuronal population from spared cortex evoked by a trimmed whisker. **(A)** Experimental timeline/design – continued from Figure 1. To induce single whisker experience (SWE), whiskers were trimmed (all contralateral whiskers but principal whisker) every two days and animals were imaged two more sessions at day ~12 and ~20 after trimming (grey shaded region). (B-C) Representative in vivo two-photon microscopy images from L2/3 neurons of the spared *w*S1 in *Syngap1^+/+^* (**B**) and *Syngap1^+/−^* mice (**C**) and representative ΔF/F traces from the same cells as in images before (baseline) and after (plasticity) SWE. Circles show low- (blue), mid- (beige) and high- (red) active cells. (**D**) Representation of the experimental design consisting in deflecting the trimmed/non-principal whisker-(NPW) and imaging in the spared/principal whisker (PW) wS1. (**E**) Change in response amplitude (the ratio of plasticity / baseline) for the trimmed/NPW stimulation for each cell class in the spared *w*S1 in *Syngap1^+/+^* (n=146 low-active cells, n=68 mid-active cells, n=16 high-active cells from 4 mice) and *Syngap1^+/−^* mice (n=194 low-active cells, n=91 mid-active cells, n=22 high-active cells from 6 mice; Mann Whitney test U=11532, p=0.0034). (**F**) Representative stimulus-evoked calcium transients of low active cells in *Syngap1^+/+^* and *Syngap1^+/−^* mice from the spared *w*S1 before (baseline) and after (plasticity) SWE. Grey area represents the stimulation window for analysis. (**G**) Bar graphs showing potentiation and depression of all three populations after SWE in *Syngap1^+/+^* (n=146 low-active cells, n=68 mid-active cells, n=16 high-active cells from 4 mice) and *Syngap1^+/−^* mice (n=194 low-active cells, n=91 mid-active cells, n=22 high-active cells from 6 mice; Mann Whitney test U=30073, p=0.0033). Medians are represented by red or black dashed lines. Circles are individual cell values. **p ≤ 0.01.

Impaired plasticity within the low-active population in *Syngap1*^+/−^ mice was not restricted to only responses evoked during NPW deflections in the spared area of *w*S1. We more broadly observed reduced plasticity of the “silent” neuronal population in *w*S1 from areas expected to scale up overall activity after SWE. For example, reduced plasticity in the low-active population was observed in the spared cortical area evoked from the untrimmed/principal whisker (PW; Fig. 3A-C), as well as in a deprived cortical area evoked with the untrimmed/NPW (Fig. 3D-F). In this latter experiment, we also observed a genotype effect on the mid-active neurons (Fig. 3E), which reduced their activity to a greater extent in *Syngap1*^+/−^ mice compared to littermate controls. This effect, along with poor scaling up of the low-active population, combined with a normal scaling-down of high-active neurons in *Syngap1*^+/−^ animals, drove an overall reduction in spared/NPW-driven neuronal responses in the deprived area (Fig. 3F). These data indicate that the process of surround potentiation (Gambino and Holtmaat, 2012), which reflects the strengthening of spared whisker inputs within neighboring receptive areas, is severely disrupted in *Syngap1* mice.

**Figure 3.**
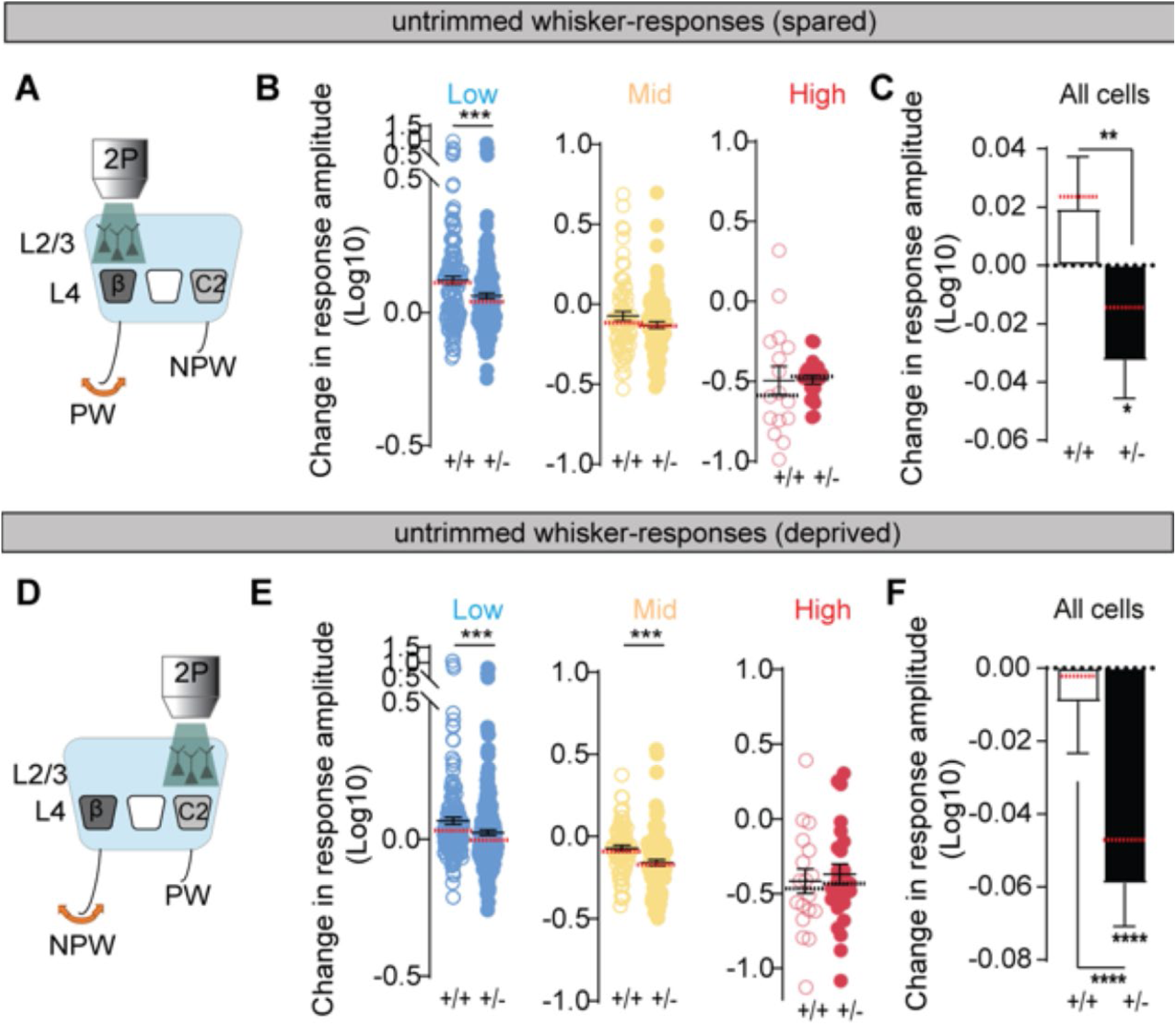
Plasticity of neuronal population responses evoked by the spared whisker in spared and deprived cortex is reduced in *Syngap1* mice. (**A**) Representation of the experimental design consisting in deflecting the untrimmed/principal whisker (PW) and imaging in the spared *w*S1. (**B**) Change in PW-response amplitude after SWE for each cell class in the spared wS1of *Syngap1^+/+^* (n= 145 low-active cells, 69 mid-active cells, 16 high-active cells from 4 mice; Mann Whitney test U= 10837, p= 0.0002) and *Syngap1^+/−^* mice (n =195 low-active cells, 91 mid-active cells, 21 high-active cells from 6 mice). (**C**) Change in PW-response amplitude after SWE for cells in the spared *w*S1 in *Syngap1^+/+^* (n= 230 neurons from 4 mice) and *Syngap1^+/−^* mice (n= 307 neurons from 6 mice; Wilcoxon Signed Tank Test, p= 0.0279; Mann Whitney test U=29680, p= 0.0016). (**D**) Representation of the experimental design consisting in deflecting the untrimmed/non-principal whisker-(NPW) and imaging in the deprived wS1. (**E**) Change in response amplitude after SWE for each cell class in the deprived *w*S1 of *Syngap1^+/+^* (n= 160 low-active cells, 75 mid-active cells, 18 high-active cells from 4 mice) and *Syngap1^+/−^* mice (n = 220 low-active cells, 102 mid-active cells, 25 high-active cells from 6 mice; Mann Whitney test U= 13849, p= 0.0004 for low-active cell comparison; Mann Whitney test U= 2535, p= 0.0001 for mid-active cell comparison). (**F**) Change in NPW-response amplitude after SWE for all cells in the deprived wS1 in *Syngap1^+/+^* (n= 253 neurons from 4 mice) and *Syngap1^+/−^* mice (n= 347 neurons from 6 mice; Wilcoxon Signed Tank Test, p<0.0001; Mann Whitney test U=35646, p< 0.0001). Bars represent mean ± SEM. Medians are represented by red or black dashed lines. Circles are individual cell values. *p ≤ 0.05, **p ≤ 0.01, ***p ≤ 0.001 and ****p ≤ 0.0001.

We next measured population plasticity in a deprived cortical area evoked from its trimmed/PW (Fig. S1A). Interestingly, changes in a deprived area during PW stimulation were not different between genotypes in any neuronal subpopulation (Figure S1B-D). This highlights the specific requirement of *Syngap1* to facilitate potentiation of spared whisker inputs, both within the principal receptive area and those that surround it. Finally, we examined whether SWE altered the IOS imaging area of spared and deprived areas of *w*S1 by deflecting the PW (Fig.S2). IOS imaging before and after whisker trimming revealed a significant whisker map expansion of the spared area regardless of initial area of the signal, though there was no difference between genotypes (Figure. S2A-C). The deprived whisker map remained unaltered by whisker trimming, regardless of genotype and initial area of the signal (Fig. S2, D-F).

### Syngap1 is required for experience-dependent reorganization of presynaptic inputs within L2/3 wS1

Potentiation of spared whisker inputs is thought to involve synapse strengthening mechanisms (Feldman, 2009; Gambino and Holtmaat, 2012). Therefore, *Syngap1* may regulate the potentiation of subpopulation response activity driven by spared whisker inputs by promoting synapse strengthening mechanisms. To test this idea *in vivo*, *Syngap1*^+/−^ mice were crossed with Thy1-eGFP mice (Feng et al., 2000) to enable repeated imaging of the same axons in L2/3 wS1 before and after SWE (Fig. 4A, B). IOS imaging was employed to identify the β whisker receptive area within *w*S1 of each mouse (Fig. 4B). Two-photon imaging was performed within L2/3 of the β receptive field during baseline sessions (no trimming) and sessions occurring after the initiation of SWE (Fig. 4A, B). As previously described for L1 in WT mice (De Paola et al., 2006; Qiao et al., 2016), most axonal boutons visualized in L2/3 were stable during baseline sessions and there was no effect of genotype on bouton stability during this period (Fig. 4C, D). As expected, SWE increased the turnover rate (TOR) of synaptic boutons in *Syngap1*^+/+^ mice (Fig. 4C, E). In contrast, TOR *decreased* in *Syngap1*^+/−^ mice after SWE, which resulted in a large genotype effect in this measure during trimming sessions (Fig. 4C, E). The genotype effect for TOR was largely driven by alterations to SWE-induced synaptic bouton formation in *Syngap1*^+/−^ mice. While bouton formation rate increased in *Syngap1*^+/+^ mice in response to SWE, it *decreased* in *Syngap1*^+/−^ mice (Fig. 4F), resulting in a strong genotype effect in this measure. In contrast, there was no change in the elimination rate in either genotype in response to SWE (Fig. 4G). However, when comparing genotypes, *Syngap1*^+/−^ mice exhibited a slight, but significant, reduction in SWE-induced changes in bouton elimination rates compared to *Syngap1*^+/+^ mice (Fig. 4F). This small genotype effect was most likely explained by opposing non-significant trends within each group. Taken together, these data demonstrate that *Syngap1* is required for experience-dependent structural plasticity of presynaptic inputs in sensory cortex.

**Figure 4.**
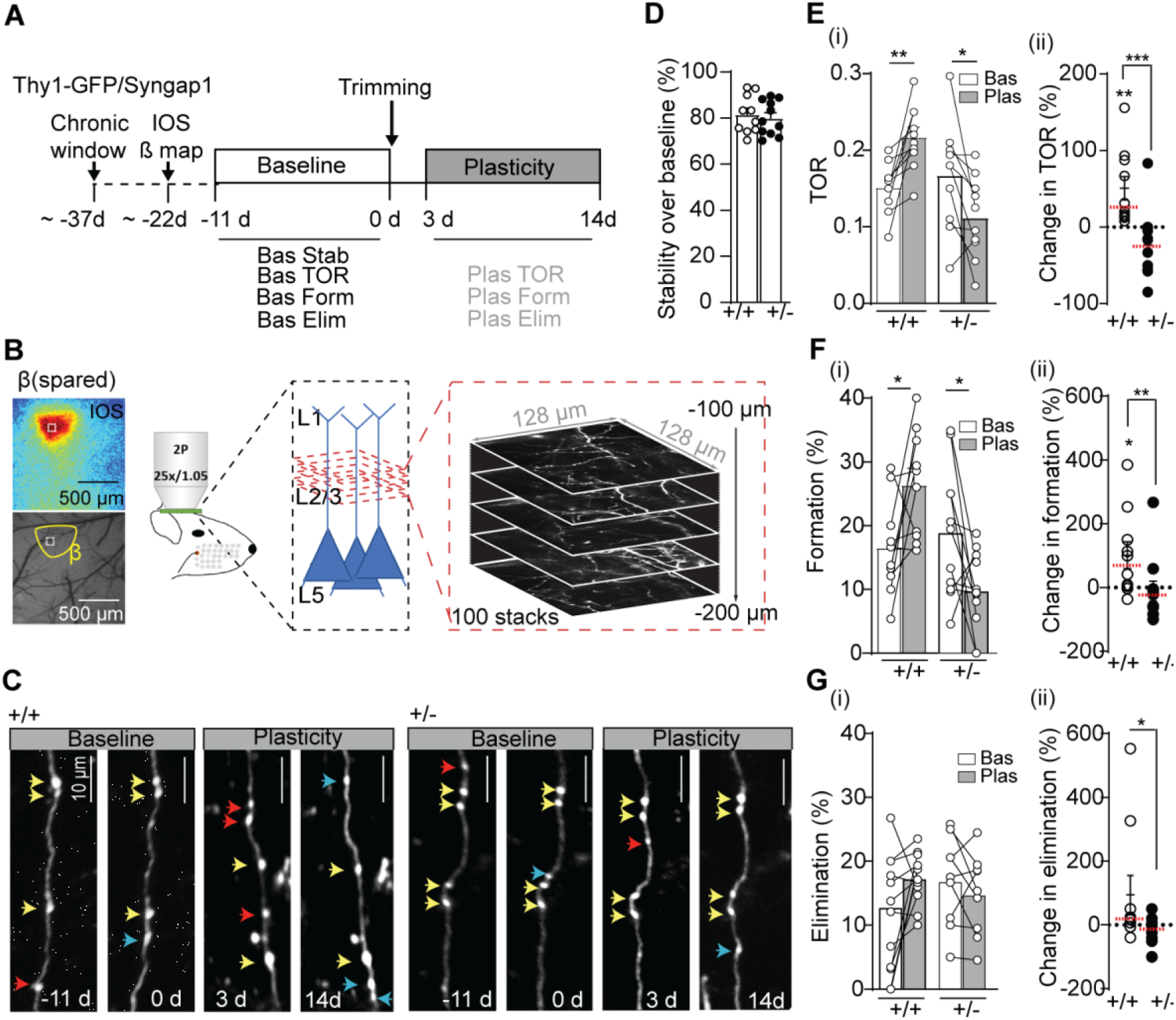
*Syngap1* heterozygosity disrupts formation of SWE-driven wS1 synaptic bouton dynamics. (**A**) Experimental timeline. Thy1-GFP/*Syngap1^+/+^* or Thy1-GFP/*Syngap1^+/−^* mice were implanted with a chronic cranial window and recovered for at least two weeks before Intrinsic Optical Signal (IOS) imaging. IOS was used to identify the cortical location activated by the deflection of the β whisker. After two baseline two-photon sessions were performed, whiskers were trimmed (all contralateral whiskers but β) every two days and animals were imaged for two more sessions. (**B**) Representation of in vivo imaging procedure of axonal bouton segments in wS1 L2/3 in Thy1-GFP/*Syngap1* mice. (**C**) Images obtained in different baseline and plasticity sessions showing axonal dynamics in *Syngap1^+/+^* and *Syngap1^+/−^* mice. Yellow arrows indicate stable axonal boutons, blue arrows indicate new boutons and red arrows indicate lost boutons. (**D**) Bouton stability over baseline sessions in *Syngap1^+/+^* and *Syngap1^+/−^* mice. (**E**) (i) Axonal bouton turnover rate (TOR) before and after single whisker experience (SWE) in *Syngap1^+/+^* (paired t test, t_9_=4.08, p= 0.0028; n= 10 mice) and *Syngap1^+/−^* mice (paired t test, t_9_=2.735, p= 0.0230; n= 10 mice). (ii) Change in TOR in *Syngap1^+/+^* (one sample t test, t_9_=3.272, p= 0.0096; n= 10 mice) and *Syngap1^+/−^* mice (Wilcoxon Signed Tank Test, p= 0.0977; n= 10 mice; Mann Whitney test U=7.00, p= 0.0005). (**F**) (i) Percentage of axonal bouton formation before and after SWE in *Syngap1^+/+^* (paired t test t_9_=2.59, p= 0.0294; n= 10 mice) and *Syngap1^+/−^* mice (paired t test t_10_=2.321, p= 0.0427; n= 11 mice). (ii) Change in axonal bouton formation in *Syngap1^+/+^* (one Sample t test t_9_=2.42, p= 0.0383; n= 10 mice) and *Syngap1^+/−^* mice (n= 11 mice; Mann Whitney test U=18.00, p= 0.0078). (**G**) (i) Percentage of axonal bouton elimination before and after SWE in *Syngap1^+/+^* (n= 10 mice) and *Syngap1^+/−^* mice (n= 10 mice). (ii) Change in axonal bouton elimination in *Syngap1^+/+^* (n= 10 mice) and *Syngap1^+/−^* mice (n= 10 mice; Mann Whitney test U=23.00, p= 0.0433). Data obtained from a total of 549 boutons in *Syngap1^+/+^* mice and 478 boutons in *Syngap1^+/−^* mice (n=11). Bars represent mean or mean ± SEM. Medians are represented by red or black dashed lines. Circles are animal means. *p ≤ 0.05, **p ≤ 0.01 and ***p ≤ 0.001.

### Syngap1 is required for spike-timing-dependent potentiation of synaptic inputs in L2/3 wS1 neurons

Hebbian plasticity of excitatory synapses in upper lamina of wS1 is believed to be a crucial physiological mechanism for cortical map plasticity during whisker experience (Allen et al., 2003; Celikel et al., 2004; Feldman, 2000; Feldman and Brecht, 2005; Feldman et al., 1999). Moreover, the potentiation of spared whisker inputs in surrounding deprived areas occurs through spike-timing-dependent plasticity (Gambino and Holtmaat, 2012). Given that a form of surround potentiation is disrupted in *Syngap1*^+/−^ mice (Fig. 3E-F), we reasoned that spike-timing dependent plasticity may also be impaired in these animals. Using thalamocortical slices containing wS1, we induced spike-timing-dependent plasticity in synapses from L4 to L2/3 by precisely pairing L4 electrical stimulation with L2/3 neuron spiking (Fig. 5A). Spike-timing-dependent long-term potentiation (STD-LTP) in L2/3 neurons was reliably induced in slices produced from *Syngap1*^+/+^ animals, but not in slices produced from *Syngap1*^+/−^ littermates (Fig. 5B-D). The role of *Syngap1* in spike timing-dependent-plasticity was restricted to potentiation because STD long-term depression was not different between genotypes (Fig. 5E-F). Control experiments ruled out any possibility of running down of stimulation in *Syngap1*^+/−^ mice because plasticity was not observed in any genotype when EPSPs were elicited without postsynaptic spikes during the pairing period (Fig. 5G-H).

**Figure 5.**
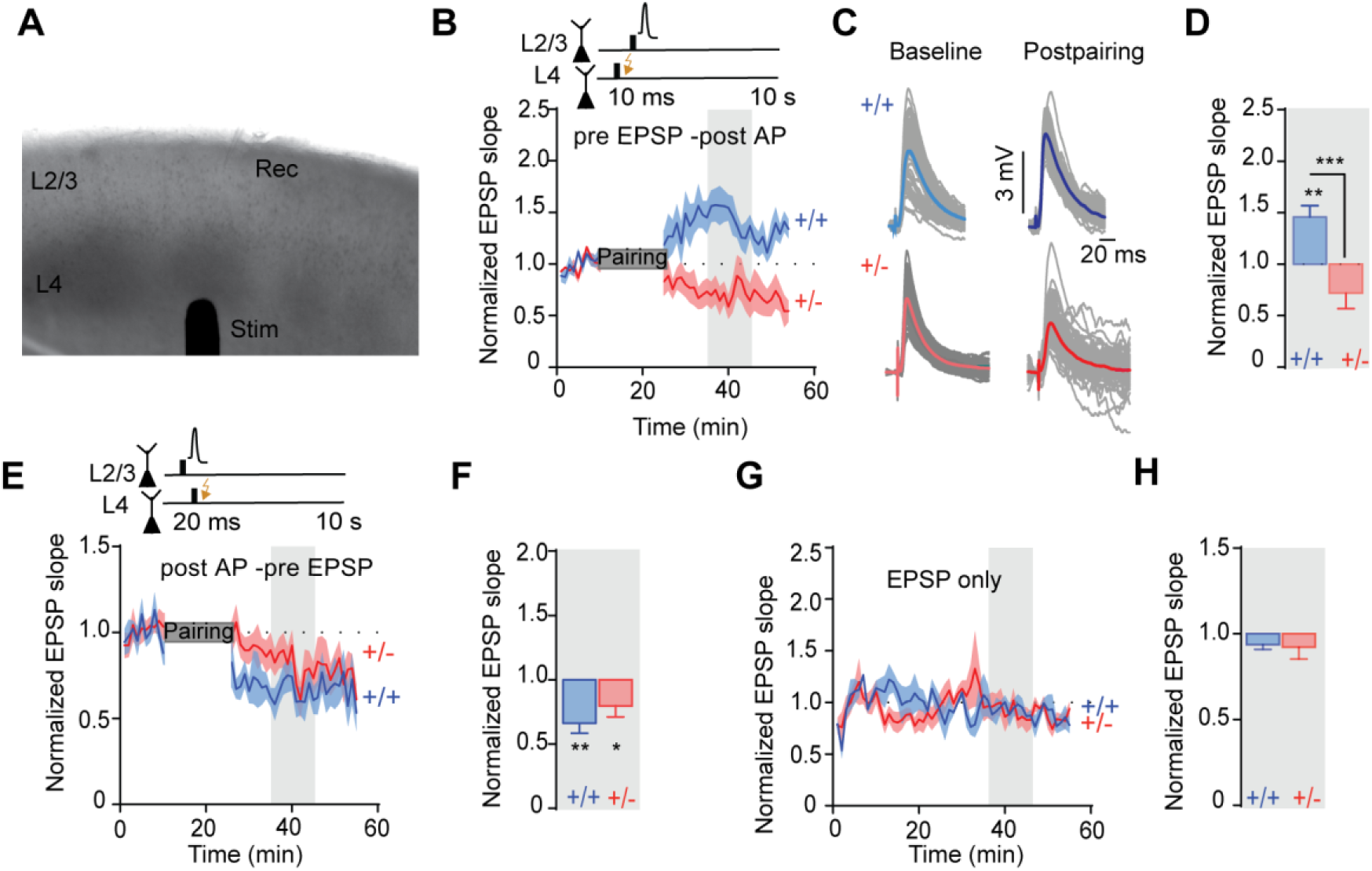
*Syngap1* is required for spike timing dependent long-term potentiation in Layer 2/3 neurons of wS1. **(A)** Image showing L4 barrels, position of stimulation electrode and recording electrode in wS1-containing thalamocortical slice. **(B)** Induction of spike-timing dependent long-term potentiation (STD-LTP) in wS1 L2/3 neurons in *Syngap1^+/+^* (n=10 cells from 8 mice) and *Syngap1^+/−^* mice (n=9 from 6 mice). The shaded area represents the time points where the quantification was performed. **(C)** Representative traces showing excitatory postsynaptic potentials (EPSPs) before (baseline) and after (plasticity) the induction of STD-LTP in *Syngap1^+/+^* and *Syngap1^+/−^* mice. **(D)** Change in EPSP slope after STD-LTP induction in *Syngap1^+/+^* (n=10 cells from 8 mice; one sample t-test t_9_=4.048, p=0.0029) and *Syngap1^+/−^* mice (n=8 cells from 6 mice; One sample t test t_7_=2.307, p= 0.0544; Unpaired t-test, t16=4.41, p= 0.0004). **(E)** Induction of spike-timing dependent long-term depression (STD-LTD) in wS1 L2/3 neurons in *Syngap1^+/+^* (n=9 neurons from 5 mice) and *Syngap1^+/−^* mice (n= 12 neurons from 8 mice). The shaded area represents the time points where the quantification was performed. **(F)** Change in EPSP slope after STD-LTD induction in *Syngap1^+/+^* (n= 9 cells from 5 mice; One sample t-test, t8=4.375, p=0.0024) and *Syngap1^+/−^* mice (n=12 neurons from 8 mice, One sample t-test t_11_=2.418, p=0.0341). **(G)** Control experiments in which only EPSP were evoked during the pairing period (n=3 *Syngap1^+/+^* neurons from 3 mice, n=4 *Syngap1^+/−^* mice neurons from 2 mice) The shaded area represents the time points where the quantification was performed. **(H)** Change in EPSP slope in control experiments (n=3 *Syngap1^+/+^* neurons from 3 mice, n=4 *Syngap1^+/−^* neurons from 2 mice). Bars represent mean ± SEM. *p ≤ 0.05, **p ≤ 0.01 and ***p ≤ 0.001.

### Syngap1-mediated regulation of L2/3 subpopulation response plasticity is not due to altered wS1 development

We hypothesized that *Syngap1* regulates experience-dependent changes in cortical population activity by initiating dynamic cellular processes that promote synaptic strengthening. However, it is also possible that alterations in the development of SSC circuits may contribute to these observations (Michaelson et al., 2018). We reasoned that if SynGAP protein controlled dynamic cellular processes, such as experience-dependent synapse strengthening, then re-expressing this protein in adult *Syngap1* mutant animals should improve neuronal population plasticity driven by SWE. To test this, we performed repeated GCaMP6s imaging in the same neuronal populations from *Syngap1^+/lx-st^*/Cre-ER/Thy1-GCaMP6s mice treated with either vehicle or tamoxifen. We have extensively validated the *Syngap1^+/lx-st^*/Cre-ER model in previous studies (Aceti et al., 2015; Clement et al., 2012; Creson et al., 2019; Ozkan et al., 2014). These animals are haploinsufficient for *Syngap1* (e.g. express half the expected SynGAP protein) and tamoxifen injections reactivate the disrupted *Syngap1* allele leading to restoration of SynGAP protein to endogenous levels. The overall design of this study was similar to that of Fig. 1A, except that *Syngap1^+/lx-st^*/Cre-ER/Thy1-GCaMP6s mice were used and injected with either vehicle or tamoxifen at PND60 (Fig. 6A). Importantly, tamoxifen treatments significantly increased SynGAP protein expression (Fig. S3A). After PND90, mice were subjected to SWE and serial two-photon imaging (Fig. S3B; Fig. 6A). During baseline imaging sessions (Fig. S3B), responses from the entire neuronal population were stable, with no overall differences between genotypes (Fig. S3C). Neurons were then categorized into the three subpopulations based on baseline response amplitude. The low- and mid-activity populations were also stable, with no differences between *Syngap1* mutants (Vehicle-treated mice) and rescue animals (TMX-treated mice; Fig. S3C-D). However, in the high-activity population, there was a slight increase in baseline activity of neurons in *Syngap1* rescue animals compared to controls (Fig.S3C-D). During SWE, we observed a lack of experience-dependent upregulation of GCaMP6s responses in the low-active (i.e. silent) population in vehicle-treated *Syngap1* mutant mice (Fig. 6B, D-E; Fig. S4A-B). Down-regulation of the high-active population occurred normally in these mice (Fig. 6B, D-E). Because Syngap1+/lx-st animals are haploinsufficient for *Syngap1*, the findings in the “silent” population are consistent with observations made previously in conventional *Syngap1*^+/-^/ Thy1-GCaMP6s mice (Figs. 2-3; Fig. S1), demonstrating impaired sensory-driven neuronal potentiation of the “silent” population in two distinct *Syngap1* mutant strains. Strikingly, re-expressing the *Syngap1* gene in *Syngap1*^+/lx-st^/Cre-ER/Thy1-GCaMP6s mice by injecting tamoxifen (Fig. S3A) restored response plasticity within the low-active “silent” population during NPW stimulations (Fig. 6C, D-E). Restoration of low-active “silent” population plasticity was also observed when sensory responses were evoked by the PW (Fig. S4A-B). The rescue of “silent” population plasticity in tamoxifen-treated mice had a significant impact on the overall cortical population response dynamics during SWE. We observed clear shifts toward potentiation of response dynamics of the combined population during SWE when comparing vehicle- and tamoxifen-treated animals (Fig. 6F-G; Fig. S4C-D).

**Figure 6.**
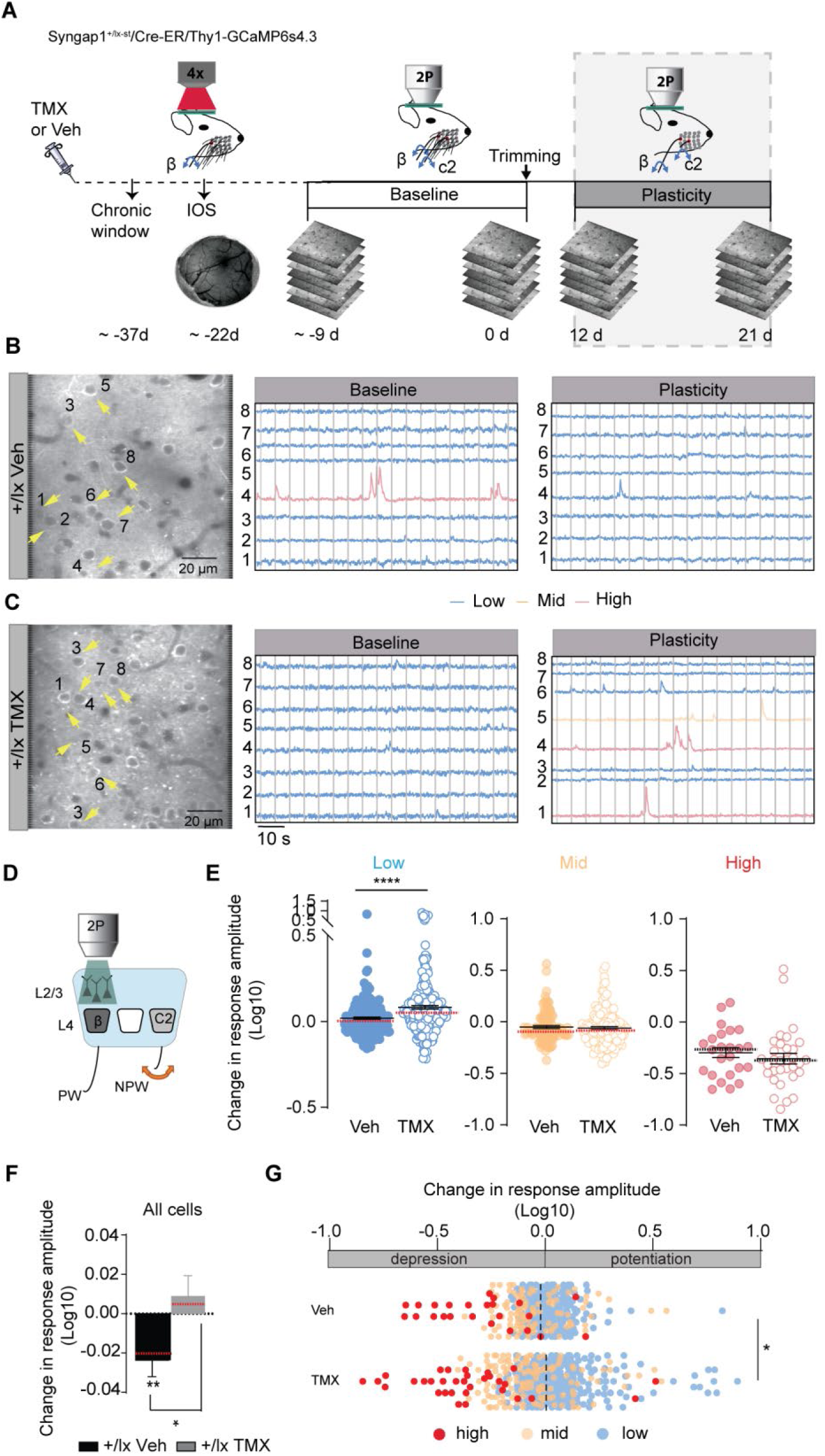
Adult re-expression of *Syngap1* reactivates experience-dependent cortical population plasticity. (**A**) Experimental timeline. *Syngap1^+/lx-st^*/Cre-ER/Thy1-GCaMP6s mice were IP injected with tamoxifen (TMX) or vehicle (Veh) and 30 days later were implanted with a chronic cranial window and allowed to recover for at least two weeks before Intrinsic Optical Signal (IOS) imaging. IOS was used to identify the cortical locations activated by the deflection of the principal whisker (PW). Whiskers were trimmed (all but the PW) every two days and animals were imaged in the spared area of *w*S1 over two additional sessions (grey shaded region). (B-C) Representative in vivo two-photon microscopy images from L2/3 neurons of the spared area of wS1 in Veh-treated (**B**) and TMX-treated mice (**C**) and representative ΔF/F traces from the same cells as in images before (baseline) and after single whisker experience (SWE). Arrows indicate cells with traces shown on the right. (**D**) Representation of the experimental design consisting of deflecting the trimmed/NPW and imaging in the spared barrel region. (**E**) Change in trimmed/NPW-response amplitude after SWE for each cell class in the spared *w*S1 of Veh-treated (n=218 low-active cells, 102 mid-active cells, 24 high-active cells from 5 mice) and TMX-treated mice (n=283 low-active cells, 132 mid-active cells, 32 high-active cells from 6 mice; Mann Whitney test U= 23516, p<0.0001). (**F**) Change in trimmed/NPW-response amplitude of all cells after SWE for the spared wS1 in Veh-treated (n= 344 neurons from 5 mice; Wilcoxon Signed Tank Test, p=0.0033) and TMX-treated mice (n= 447 neurons from 6 mice; Mann Whitney test U=68786, p= 0.011). (**G**) Bar graphs showing potentiation and depression of all three populations after SWE in Veh-treated (n=218 low-active cells, 102 mid-active cells, 24 high-active cells from 5 mice) and TMX-treated mice (n=283 low-active cells, 132 mid-active cells, 32 high-active cells from 6 mice; Mann Whitney test U=68787, p=0.0110). Bars represent mean ± SEM. Medians are represented by red or black dashed lines. Circles are individual cell values. *p ≤ 0.05, **p ≤ 0.01, ***p ≤ 0.001 and ****p ≤ 0.0001.

## Discussion

The reorganization of synapse structure and function is a fine-scale substrate that underlies changes in cortical population activity in response to sensory experience (Buonomano and Merzenich, 1998; Darian-Smith and Gilbert, 1994; Margolis et al., 2012). However, to our knowledge, opposing functional changes within defined neuronal subpopulations of the same local cortical network during sensory experience has only been described in a single publication (Margolis et al., 2012). Importantly, we confirmed many of the original findings from the Margolis *et al.* study even though we used a distinct genetically encoded calcium sensor (GCaMP6s vs. YC3.60) expressed using a different method (stably expressed transgene in the germline vs. direct viral delivery to cortex). L2/3 neurons in wS1 were classified into three subpopulations based on response amplitude and each of these populations exhibited stable levels of activity during baseline imaging sessions (i.e. no whisker trimming). This finding was critical because the analysis of population-based plasticity during SWE relies on the stability of these populations during baseline conditions. Most importantly, we replicated the key finding of the Margolis *et al* study; the observation that spatially intermingled cells within wS1 exhibited highly specific and sometimes opposing changes in activity during SWE (Margolis et al., 2012). We observed opposing changes in responsiveness of different types of functionally defined cortical neurons within the same imaging fields in wS1. Neurons defined as low active, or “silent”, increased activity in response to SWE, while high-active neurons exhibited reduced sensory responsiveness during trimming sessions. Margolis *et al.* argued that the differences observed in these subpopulations emerged from their distinct role within the barrel cortex network. Low-active neurons, which are the vast majority of neurons in wS1, weakly respond to whisker stimulation, but exhibit the largest change in response to SWE. Their ability to extensively scale up activity is thought to enable remapping of cortical circuits in response to experience, which would change computational functions within the network. In this context, scaling down of spatially intermingled high-active neurons may be a compensatory response within the network to maintain overall firing rates in the face of massively upregulated synaptic inputs in the low-active population. However, scaling down of a small population of neurons could be explained in other ways, such as neuronal toxicity known to be caused by viral expression of calcium sensors, or high-active neurons could represent a distinct neuronal type, such as fast-spiking interneurons. Our studies argue against these two alternative possibilities. Neuronal toxicity in the GP4.3 transgenic mouse line has been documented to be minimal (Dana et al., 2014). Indeed, stable neuronal responses have been reported over the course of many months in this mouse line. In our experiments, we observed very few neurons with nuclear signal, even after three months of experimentation with up to six repeated imaging sessions in the same area. In addition, within the cortex, GCaMP6 is not expressed in GABAergic neurons (Dana et al., 2014). Therefore, both low- and high-active populations were comprised exclusively of glutamatergic neurons, demonstrating that neighboring neurons of the same general type can exhibit opposing changes in neuronal responses during SWE.

Our findings in *Syngap1* mutant models support the idea that distinct molecular and cellular mechanisms underlie subpopulation specific changes to neuronal activity during SWE. In addition, our results indicate that *Syngap1* is required to initiate synapse strengthening mechanisms required for experience-dependent changes in neuronal population responses. We found that SWE-mediated scaling-up of the low-active “silent” wS1 neuronal subpopulation was impaired in two distinct models of *Syngap1* haploinsufficiency. However, plasticity of high-active neurons, which scale-down their activity during whisker experience in WT mice, was unimpaired in these mice. Thus, expression of this gene promotes neuron-specific plasticity within whisker-responsive SSC networks. Several lines of evidence indicate that *Syngap1* promotes scaling-up of low-active neurons through the regulation of cellular mechanisms that strengthen synaptic connections in Layer 2/3. Similar to past reports (De Paola et al., 2006; Qiao et al., 2016), we found that whisker experience triggered axonal dynamics resulting in a net gain of new synaptic boutons in upper lamina SSC of WT mice. Thy1-eGFP mice used in this study express the fluorescent reporter exclusively within excitatory neurons (Feng et al., 2000), demonstrating that there was a net gain of excitatory input in L2/3 in response to SWE. This is a plausible mechanism contributing to scaling up of the low-active population during whisker experience. In *Syngap1* mutant mice, however, SWE caused a reduction, rather than an increase, in the formation of new boutons. Thus, losing one copy of *Syngap1* prevented a form of experience-dependent structural plasticity that drives a net-gain of excitation within upper lamina wS1 circuits.

In addition to impaired structural plasticity, we observed impaired functional plasticity of excitatory inputs onto L2/3 wS1 neurons in these mice. Spike-timing dependent LTP in acute brain slices obtained from *Syngap1* mutants was absent in wS1 L2/3 neurons. This form of LTP is thought to contribute to whisker map expansion of the spared whiskers (Feldman, 2009; Gambino and Holtmaat, 2012). Importantly, we found that low-active neurons in the deprived area of wS1 from *Syngap1* mice failed to scale-up activity in response to stimulation of the spared whisker, a finding consistent with impaired STD-LTP. These findings in cortical circuits are consistent with known functions of SynGAP protein in hippocampal neurons. SynGAP protein is a core constituent of the postsynaptic density within dendritic spines where it dynamically regulates small GTPase signaling in an NMDA receptor and CAMKII-dependent manner to regulate AMPAR trafficking required for synapse strenthening (Gamache et al., 2020; Kilinc et al., 2018).

Finally, our data demonstrate that synaptic weakening mechanisms are intact in wS1 of *Syngap1* mice. Axonal bouton elimination rates and STD-LTD were not different between genotypes in L2/3 of wS1. This is consistent with known hippocampal functions of SynGAP, where synaptically-evoked LTD at Schaeffer collaterals in *Syngap1* mutants is unimpaired (Kim et al., 2003). The observation here that *Syngap1* heterozygosity disrupts synapse strengthening mechanisms in cortex, while leaving weakening mechanisms intact, may explain the overall weakening of combined neuronal population responses evoked from the spared whisker after SWE (Fig. 3D, G). Intact synapse weakening may also explain the lack of effects observed in population activity from the deprived area of cortex evoked from the trimmed/principal whisker (Fig. S1). Indeed, synapse weakening mechanisms are thought to predominate in deprived cortical areas (Feldman, 2009).

Restoration of population plasticity within “silent” wS1 neurons after adult *Syngap1* re-expression (Fig. 6) supports a model where SynGAP protein controls dynamic processes, initiated at the time of experience, to support the redistribution of neuronal population activity. Past studies support this interpretation. For example, impaired Schaffer collateral LTP in *Syngap1* mutant mice is completely rescued after adult re-expression of SynGAP protein (Ozkan et al., 2014). Importantly, dynamic Ras/ERK signaling initiated by theta-burst stimulation was also rescued in the adult hippocampus of *Syngap1* mutants in this same study. Consistent with these findings, adult re-expression of SynGAP protein also improves deficits in hippocampal theta rhythms in concert with systems memory consolidation of long-lasting contextual fear (Creson et al., 2019), which are processes that rely on intact hippocampal function (Colgin, 2016). Taken together, *Syngap1* also appears to have critical functions in adult cortical circuits that are required for the initiation of dynamic processes supporting experience-dependent remapping of neuronal population activity. Thus, impaired experience-dependent cortical plasticity and altered redistribution of cortical population activity described here is an outcome of *Syngap1* heterozygosity that unrelated to the known role of this gene to regulate developmental assembly of neural circuits (Aceti et al., 2015; Clement et al., 2012). As a result, boosting expression of SynGAP protein in patients with *SYNGAP1* haploinsufficiency may improve experience-dependent cortical plasticity, a fundamental neuronal process required for learning and behavioral adaptations. Furthermore, given that many other NDD risk genes also impact activity-dependent processes (Ebert and Greenberg, 2013) and synapse biology (Bourgeron, 2015), it is tempting to speculate that a common feature of these disorders is impaired fine-scale redistribution of cortical neuron activity during experience.

## Author Contributions

N.L. performed experiments, designed experiments, analyzed data, co-wrote the manuscript and edited the manuscript. T.V. analyzed data, interpreted data, and edited the manuscript. C.R. generated reagents. S.M. analyzed data and edited the manuscript. C.A.M. interpreted data and edited the manuscript. G.R. conceived project, designed experiments, interpreted data, co-wrote and edited the manuscript.

## Data and materials availability

The data that support the findings of this study are available from the corresponding author upon reasonable request.

## Materials and Methods

### Animals

All animal procedures were conducted in accordance with the National Institutes of Health *Guide for the Care and Use of Laboratory Animals* and all procedures were approved by the Scripps Institutional Animal Care and Use Committee. Males and females were used in all experiments indiscriminately. Mice were housed 4 or 5 per cage on a 12-h normal light–dark cycle. For experiments requiring chronic cranial window and headpost implantation, mice were singly housed following surgery with cardboard huts for the remainder of the study. We used inbred *Syngap1* constitutive (*Syngap1*^+/−^) (Kim et al., 2003) or *Syngap1* conditional rescue (*Syngap1^lx-st^*) mice (Clement et al., 2012). Each line is maintained by colony inbreeding on a mixed background of C57-BL6/129s. Every seventh generation, *Syngap1*^+/−^ or *Syngap1^lx-st^* mice are refreshed by crossing colony breeders into C57-BL6/129 F1 animals for one generation. Offspring from these crosses, like those used for this study, are then inbred for up to seven generations. For 2-photon microscopy experiments, *Syngap1*^+/−^ or *Syngap1^lx-st^* mice were crossed with Thy1-GCaMP6s4.3 (#024275) or Thy1-GFP (#007788) reporter lines, which were purchased from Jackson Laboratories. F1 offspring were used for these studies, except for animals generated for studies shown in Figure 6. To create animals used in Figure 6, a hemizygous inducible Cre driver line that our group has previously validated (JAX stock #004682) (Aceti et al., 2015; Clement et al., 2012; Creson et al., 2019; Ozkan et al., 2014) was crossed to a double mutant *Syngap1*+/lx/Thy1-GCaMP6s4.3 line. Only triple mutant animals were selected for experiments. Animals expressing the inducible Cre transgene were injected (intraperitoneal) with tamoxifen for five days starting at postnatal day (PND) 60. Tamoxifen (Sigma T5648, St. Louis, MO) was prepared by dissolving it into absolute ethanol (Acros/Fisher Scientific 61510-0010, Pittsburg, PA) (10% of final volume) by sonication to which corn oil was added for a final dosage of 100 mg/kg, injectable concentration of 20 mg/ml, and volume of 5 ml/kg. For all studies, the experimenter was blind to genotype at the time of data acquisition and analysis.

### Sensory Manipulation

For sensory deprivation experiments we employed the single whisker experience paradigm. All contralateral whiskers but one (usually, but not always, β whisker) were trimmed, ipsilateral whiskers were left intact. Trimming started immediately after the last baseline session by cutting whiskers to fur level using micro scissors under microscope while mice were under low anesthesia (1.5-2% isoflurane). Whiskers were re-trimmed every other day for a maximum of 21 days by lightly anesthetizing mice (1.5-2% isoflurane).

### Slice electrophysiology

#### Preparation of barrel cortex slices

Thalamocortical slices (350 um) containing the barrel cortex were prepared as described previously (Agmon and Connors, 1991). P21 to P30 mice of either sex were decapitated under isofluorane in accordance with Scripps regulation. The brain was rapidly removed in ice-cold artificial cerebrospinal fluid (aCSF) containing (in mM): 119 NaCl, 2.5 KCl, 1.3 MgSO4, 2.5 CaCl2, 1 NaH2PO4, 11 D-glucose and 26.3 NaHCO3, pH 7.4, 300-310 mOsm bubbled with 95%CO2 and 5%O2. Slices were cut on a vibrating microtome (Candem Instruments), transferred to a submersion chamber (Warner Instruments, Hamden, CT) at 32-34C for 30min and then kept at room temperature until recording (1-6h).

All recordings were made at room temperature in standard aCSF in slices containing the barrel cortex. The barrel subfield was identified by transillumination at 4X by the presence of three to five ~ 300 uM wide barrels in L4. Whole-cell patch-clamp recordings of L2/3 excitatory cells were made in one of the barrel columns under visual guidance by infrared differential interference contrast (DIC) microscopy at 40X. L2/3 excitatory cells were identified by their soma shape and their location ~ 150 uM below the L1-L2 boundary. As expected for pyramidal neurons (Agmon and Connors, 1992; Connors and Gutnick, 1990), all cells showed regular spiking responses to positive current injections. Experiments were made with borosilicate patch-pipettes (3-5 mΩ) filled with a solution containing (in mM): 116 K-gluconate, 6 KCl, 2 NaCl, 0.5 EGTA, 20 HEPES, 4 Mg-ATP, 0.3 Na-GTP and 10 Na2-phosphocreatine (pH: 7.3, 280 mOsm)

#### Whole-Cell Recording of Spike-Timing Dependent LTP and LTD

The stimulating electrode was placed accurately in a L4 barrel under visual guidance in a trans-illuminated slice. Excitatory postsynaptic potentials (EPSP) were evoked via a concentric bipolar stimulating electrode placed within the base of a barrel in L4, vertically aligned to the site of the recording. EPSPs were evoked at a constant rate of 0.1 Hz. Timing-based plasticity was induced as previously described ((Feldman, 2000)). Briefly, after a stable baseline period of 10 min, single EPSPs were paired with single action potentials (APs) evoked by the minimum current injection required to evoke an AP at precise delay before or after each AP. To induce STD-LTP, the postsynaptic AP was evoked within 10 ms after the onset of the EPSP, whereas STP-LTD was induced by evoking the postsynaptic action potential 20 ms before the onset of the EPSP. After 75 pairing sweeps current injection was suspended and EPSP slope was monitored for 30 minutes. Presynaptic stimulation intensity remained constant throughout the experiment. Series resistance was compensated. Input resistance was monitored, and cells were discarded if the resting membrane potential changed by more than 8 mV. For quantification of LTP and LTD, the ratio of postpairing slope during 10 min beginning and 10 min after the end of pairing to that in the baseline was calculated. Only the initial slope (first 2 ms) of the EPSP was analyzed.

#### Electrophysiological data acquisition and storage

All signals were amplified using Multiclamp 700B (Molecular Devices, Sunnyville, CA), filtered at 4 kHz, digitized (10 kHz), and stored on a personal computer for offline analysis. Analog to digital conversion was performed using the Digidata 1440 A system (Molecular Devices). Data acquisitions and analyses were performed using pClamp 10.2 software package (Clampex and Clampfit programs; Molecular Devices).

### Chronic cranial window implantation

For Thy1-GCaMP6s4.3 and Thy1-GFP mice experiments, both male and female mice at least 8 weeks of age were fitted with a chronic cranial window and implanted with a titanium headpost according to established procedures with minor modifications (Michaelson et al., 2018). Briefly, animals were anesthetized with isoflurane (5% induction, 1.5–2% maintenance) and IP injected with a cocktail of dexamethasone (4 mg/kg), Rimadyl (carprofen 10 mg/kg), and Enroflox (enrofloxacin 5 mg/kg). Animals were mounted on a stereotaxic frame (David Kopf Instruments, Tujunga, CA) and body temperature was maintained with a thermal regulator (Harvard Apparatus, Holliston, MA). The scalp was shaved and sterilized with alternating swabs of Betadine and 70% alcohol. A small skin flap was removed, the periosteum was gently cleared, and the skull was scraped with a scalpel. A small circular craniotomy was made over the left barrel cortex (3-mm diameter; center relative to bregma: lateral 3.5 mm; posterior 1.8 mm) using a dental drill, and the dura was left intact. Two 3-mm glass coverslips were glued onto a 5-mm glass coverslip, and the cranial window was sealed by gluing these coverslips directly to the bone (VetBond, 3 M). The titanium headpost was implanted by adhering it directly to the bone using VetBond and then dental cement (Metabond, Parkell, Edgewood, NY). Animals recovered on a warm blanket before being placed back in their home cage. Rimadyl and Enroflox was injected (5 mg/kg) for 3 consecutive days after surgery for pain management. Animals were left to recover at least 10 days before subsequent experiments.

### Intrinsic optical signal (IOS) imaging

Animals of at least 8 weeks of age were IP injected with the sedative chlorprothixene (1 µg/g) and were anesthetized with a lower dose of isoflurane (5% induction, 0.5 % maintenance) (Juavinett et al., 2017). Imaging was performed under a 4× objective on an upright microscope frame (BW51X; Olympus, Tokyo, Japan). The barrel cortex was illuminated with 630-nm LEDs mounted on the 4× objective. The images were acquired with a Zeiss Axiocam camera (Carl Zeiss Microscopy Inc., Thornwood, NY) controlled by µManager software (Open Imaging, Inc.). Acquisition rate was approximately 10 Hz. Whiskers were deflected using a piezoelectric bending actuator controlled by a linear voltage amplifier (Piezo Systems Inc., Woburn, MA). A single sinusoidal wave with a 5-ms rise and a 5-ms decay time was generated using Clampex software (Molecular Devices, Sunnyville, CA). Bending of the piezo was calibrated using a laser-based displacement device (LD1610-0.5 Micro-Epsilon, Raleigh, NC). A single whisker deflection was approximately 200 µm at 2 mm away from the whisker pad.

#### Analysis of IOS imaging

Each IOS imaging trial consisted of a 2-s baseline imaging period followed by 40 deflections at 10 Hz. We performed 50–70 trials for each whisker and averaged them using IO and VSD Signal Processor plugin in ImageJ (Harrison et al., 2009). Images taken between 1 s and 3 s after the start of the stimulus were averaged and defined as the response. IOS images were obtained by calculating the (response – baseline)/baseline value for each pixel using custom scripts written in Matlab (MathWorks, Natick, MA), according to established procedures (Chen-Bee and Frostig, 1996; Chen-Bee et al., 2000). Investigator was blind to animal genotype at the time of the analyses. Briefly, images were first filtered with a Gaussian filter. Afterwards a baseline and a response region were manually selected in the final IOS image to minimize contamination by blood vessels (Chen-Bee and Frostig, 1996). Response size was determined as the minimum value of the response region subtracted from the median of the baseline region. Image thresholding was performed in the response region to determine the area of activation. Relative thresholding values were set at 50% of the response size for each image.

### In vivo 2-photon imaging

In vivo 2-photon imaging was performed in L2/3 of the barrel cortex. For each imaging session, mice of at least 8 weeks of age were anesthetized with isoflurane (1.5-2%). Individual sessions lasted 40-90 min, after which animals were returned to their home cage (one animal per cage). Following recovery from surgery, IOS imaging was performed through the cranial window, as described above, using light (0.5–1%) isoflurane anesthesia to locate principal whisker areas (typically β and C2 whisker). Imaging was performed with a VivoScope two-photon microscope equipped with a resonant scanner (Scientifica, UK). The light source was a Mai Tai HP 100 femtosecond-pulse laser (Spectra-Physics) running at 940 nm for GCaMP and 910 nm for GFP. The objective was a 16 × water immersion lens with 0.8 NA (Nikon) for GCaMP imaging and ULTRA 25x with 1.05 NA (Olympus) for GFP imaging. Images were acquired using ScanImage 5 (http://vidriotechnologies.com). For GCaMP6 imaging images (512 × 512 pixels, 4 × zoom, 150 × 150 µm) of L2/3 cells (70–250 µm below the pia) were collected at 10 Hz. For GFP imaging of L2/3 axonal boutons from one wS1 cortical area (usually, but not always, β) image stacks were collected 100-200 µm below the pia at 1 µm intervals. Each z-image was 1024 x 1024 pixels, 4 x zoom, and was integrated over 10-20 seconds at each depth. Laser power at the sample was estimated to be <80mW for GCaMP6 experiments and <30mW for eGFP experiments. A similar number of imaging depths and same number of imaging sessions at similar depths were acquired for each animal. The head of the animals was in a fixed position across sessions to ensure consistent orientation of imaging planes. For repeated imaging, areas were reacquired using blood vessels as landmarks.

#### Analysis of GCaMP activity in the barrel cortex

Neuronal populations residing in two different *w*S1 areas (usually populations from β and C2 receptive fields) were studied during four different imaging sessions (i.e. two baseline sessions and two plasticity sessions). For Thy1-GCaMP6s4.3/*Syngap1* mice, the first baseline session was performed at day ~ −13 (−13.10 ± 2.68) and the second baseline always at day 0 right before trimming (relative to whisker trimming day, i.e. day 0). The first plasticity sessions were completed at day ~12 (12.10± 0.10) and second plasticity sessions at day~20 (19.52± 0.98) relative to trimming day. For experiments performed with Thy1-GCaMP6s4.3/Cre-ER/*Syngap1^+/lx-st^*, the first baseline sessions were performed at day ~ −9 (−8.63 ± 0.34) and the second baseline always at day 0 right before trimming. After whisker trimming, sessions were always completed at day 12 and day 21 for the first and second plasticity sessions, respectively.

Within each imaging area, all neurons that could be visualized in each of the four sessions were studied, irrespective of levels of activity or responsiveness to the deflection of the principal whisker (PW) and non-principal whisker (NPW). PW and NPW deflection responses were analyzed for each neuron visualized in both imaging areas. For each imaging session, extraction of ΔF/F of calcium images was performed in Matlab R2015b using the FluoroSNNAP15.04.08 plugin (Patel et al., 2015) with the following parameter choices. Regions of interest (ROIs) corresponding to identifiable cell bodies along the four imaging sessions were selected manually. The fluorescence time course was measured by averaging all pixels within the ROI, then corrected for neuropil contamination. The neuropil ROIs were also manually drawn where there were no visible cell bodies and were the same for all cells within an imaging frame. After neuropil correction, the ΔF/F of each ROI was calculated as (F − F0)/F0, where F0 was the mean of the lower 50% of the proceeding 10-s period. For the first 10-s period, a minimum value of F0 was used (Patel et al., 2015). A template search-based algorithm was used in order to detect calcium events using built-in templates in FluoroSNNAP15.04.08.

Whisker-stimulation-induced activity was recoded over a 2-min period from the same ROIs (PW and NPW stimulation activity was recorded for each ROI). Whisker stimulation consisted of 10 whisker stimulations at 20 Hz with an intertrain interval of 5.12 s. For each type of whisker deflection (i.e. PW or NPW) a total of 21 trains were given during a 2-min period, from which the greatest ΔF/F value per cell per type of whisker deflection (PW or NPW) was taken for analysis. We used custom-written R scripts for the extraction and processing of the maximum ΔF/F amplitude induced by PW and NPW stimulation (1 s window/stim) per session in each cell. Based on their maximum ΔF/F response to the PW stimulations during baseline sessions, cells were classified into low-, mid- and high-active cells as previously described (Margolis et al., 2012). Briefly, low-active cells comprise 63.3 % of the total population included for analyses. Mid-active and high-active represent the 29.6% and 7.1% of the total population, respectively. Next, the maximum ΔF/F response per cell were averaged per type of session (i.e. baseline or plasticity) and the change in the averaged ΔF/F (i.e. ratio of plasticity / baseline) of each cell was used to study the effect of whisker trimming on neuronal dynamics.

### Analysis of axon bouton dynamics in the barrel cortex

The exact same axon shafts residing in a specific *w*S1 area (usually, but not always β receptive area) were imaged throughout four imaging sessions (i.e. two baseline and two plasticity sessions). Relative to whisker trimming day (i.e. day 0), the first baseline was always collected at day – 11 and the second baseline always at day 0, right before whisker trimming (all contralateral whiskers but β, typically). Plasticity sessions were always completed at day 3 and day 14 relative to trimming day. Axonal segments (20-125 µm) visible in L2/3 of the β receptive area were identified from the image stacks from their location and morphology. Three to eleven axonal segments per animal were collected and considered for analysis if visible throughout the four imaging sessions. Within animals, axon bouton dynamics were compared between baseline and plasticity sessions (eleven-day time window in both types of sessions). For calculation of bouton turnover rate (TOR), formation and elimination rates, z-stack images containing the axon segment of interest were sum intensity projected to include the whole arbor. Next, the axon of interest was traced manually with a line selection tool in ImageJ (National Institutes of Health). The intensity along the selected line was measured. Boutons that exhibited a peak intensity > 1.5 times the average axon shaft intensity on session1 were selected for analysis. A bouton was scored as stable if this bouton maintained its peak intensity >1.5 times than the axon shaft. New peaks of intensity >1.5 times the average axon segment intensity was scored as gain. If the peak of intensity of a bouton dropped below 1.2 times the average axon segment intensity, it was scored as a loss. In the case of finding intensity peaks in close proximity to each other, intensity peaks were scored as distinct boutons if they were at least 2 µm apart. TOR was calculated as TOR (t_1_, t_2_) = (N_gained_ + N_lost_)/(2 × N(t_1_) (De Paola et al., 2006; Grillo et al., 2013). The percentage of boutons formed or eliminated is defined as the number of boutons formed or eliminated divided by the number of existing boutons at the first session compared. The change in TOR, change in formation and change in elimination refers to the percentage of increment measured over a given interval (Pan et al., 2010).

### Statistics

For statistical analysis applied to specific comparisons see **Table S1**. Data analyses were conducted in MATLAB (MathWorks, version 2013b and 2015b, Natick, MA), R studio (RStudio, Inc) and GraphPad Prism 8 (GraphPad Software, CA). D’Agostino–Pearson omnibus normality tests were applied to determine data distributions, and the appropriate parametric or nonparametric statistical test was performed accordingly. For analysis of electrophysiological data, one sample t-test, two-sided Student’s t test and two-way ANOVA (mixed model) was used. For analysis of axon boutons imaging data, the following tests were used: two-sided paired Student’s t test was used for within genotype comparisons of TOR, formation and elimination before and after sensory deprivation. For within genotype comparisons of change in TOR, change in formation and change in elimination one sample t-test and Wilcoxon Signed Rank tests were used. For genotype comparisons of change in TOR, change in formation and change in elimination Mann-Whitney U tests were used. For within genotype comparisons of GCaMP imaging data one sample t-test and Wilcoxon Signed Rank tests were used. For genotype comparisons of GCaMP imaging data two-sided Student’s t test or Mann–Whitney U test were used. Data throughout the text are presented as mean ± SEM. Differences were considered to be significant for *p* < 0.05.

---

**Figure S1.**
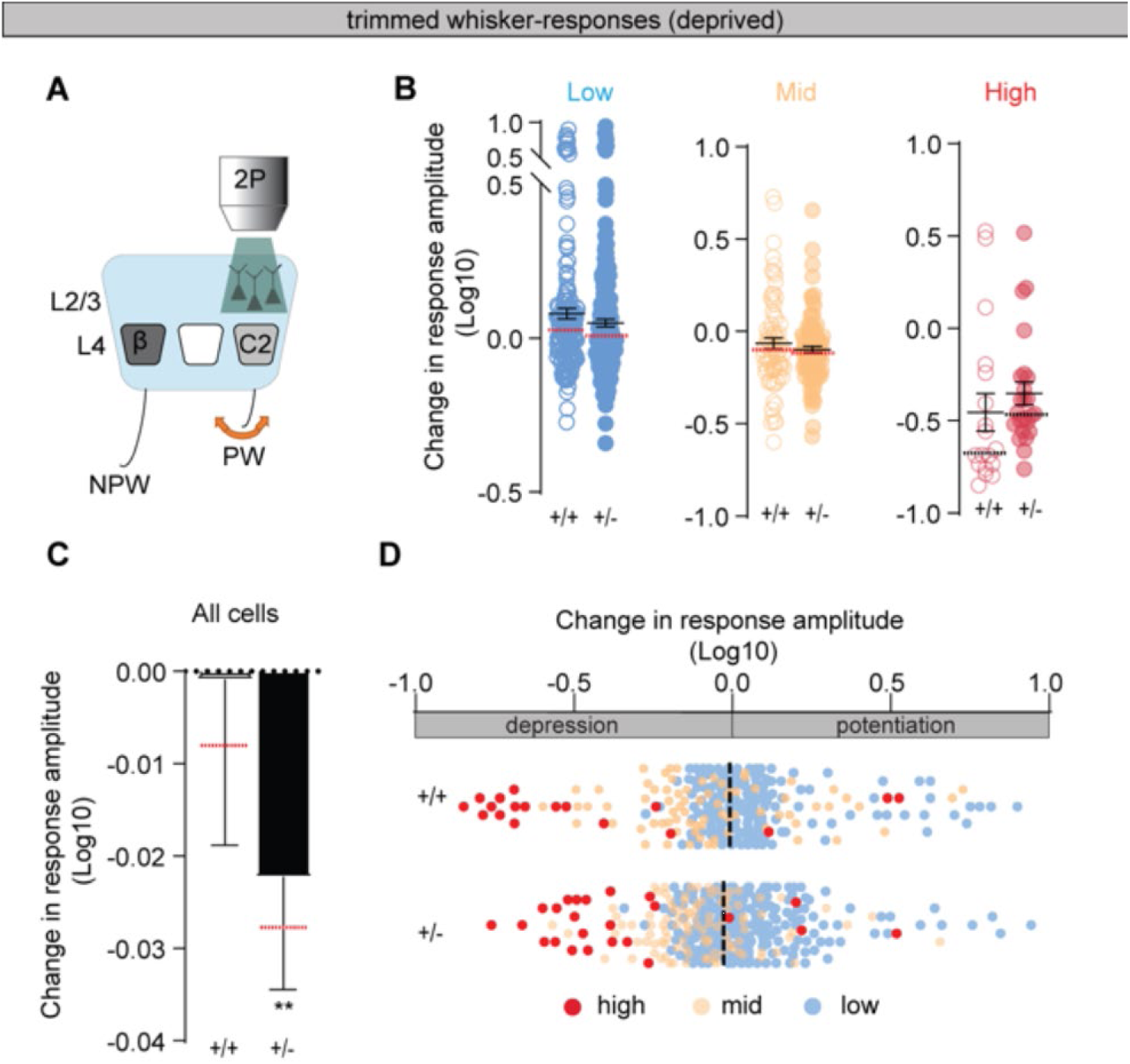
Plasticity of neuronal population activity in deprived wS1 evoked from the trimmed whisker is not altered in *Syngap1* mice. (**A**) Representation of the experimental design consisting of deflecting the trimmed/principal whisker-(PW) and imaging in the deprived whisker primary somatosensory cortex *w*S1. (**B**) Change in response amplitude after SWE for the trimmed PW-stimulation for each cell class in the deprived wS1 in *Syngap1^+/+^* (n=160 low-active cells, n=75 mid-active cells, n=18 high-active cells from 4 mice) and *Syngap1^+/−^* mice (n=220 low-active cells, n=103 mid-active cells, n=24 high-active cells from 6 mice). (**C**) Change in trimmed/PW-response amplitude of all cells after single whisker experience (SWE) for L2/3 neurons in the spared wS1 in *Syngap1^+/+^* (n= 253 neurons from 4 mice) and *Syngap1^+/−^* mice (n= 347 neurons from 6 mice; Wilcoxon Signed Tank Test, p=0.0027). (**D**) Bar graphs showing potentiation and depression of all three populations after SWE in *Syngap1^+/+^* (n=160 low-active cells, n=75 mid-active cells, n=18 high-active cells from 4 mice) and *Syngap1^+/−^* mice (n=220 low-active cells, n=103 mid-active cells, n=24 high-active cells from 6 mice). Bars represent mean ± SEM. Medians are represented by red or black dashed lines Circles are individual cell values. *p ≤ 0.05.

**Fig S2.**
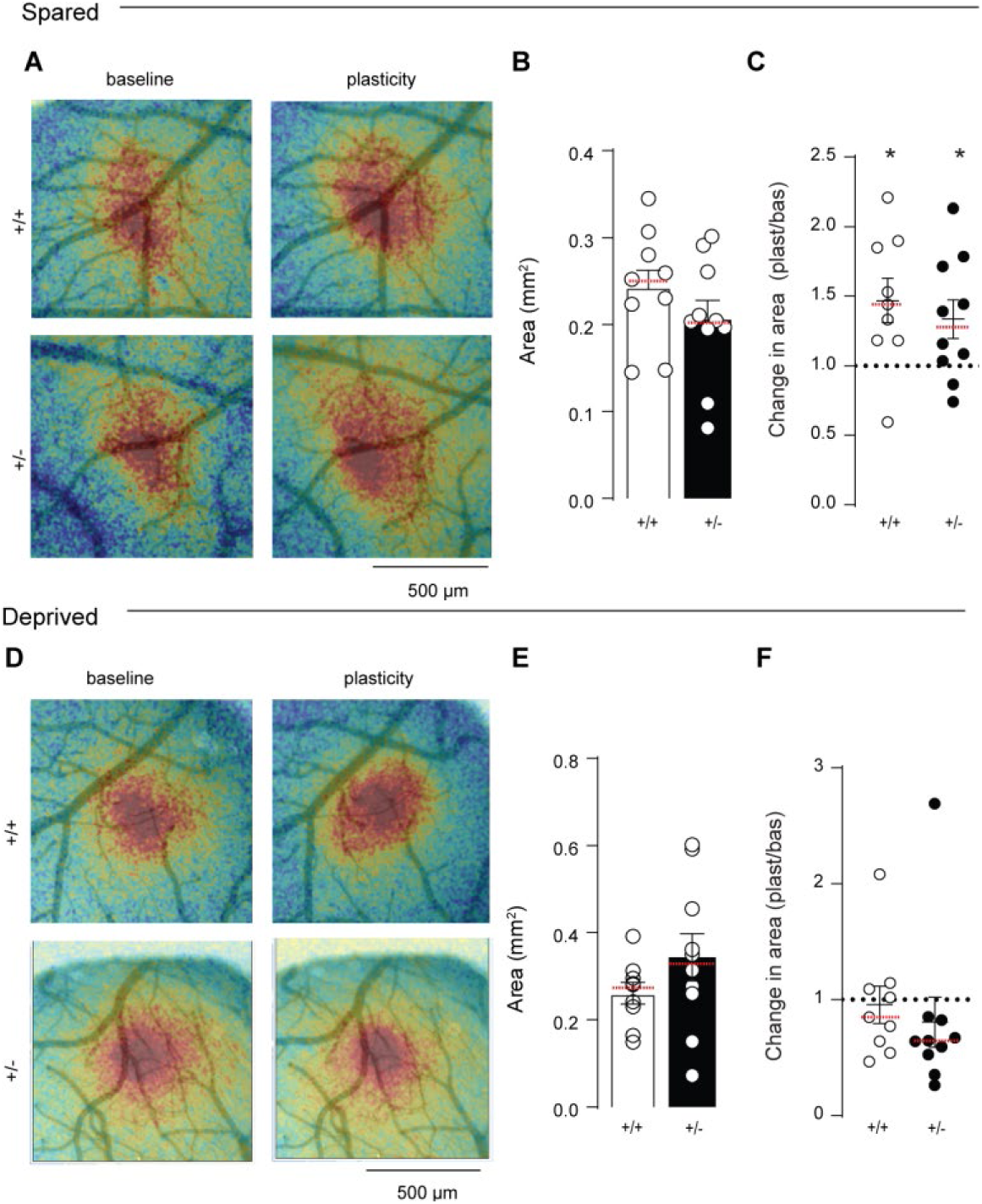
Whisker map plasticity in *Syngap1* mice measured by intrinsic optical signal imaging. (**A**) Intrinsic optical signal (IOS) images superimposed on corresponding blood vessels images showing map expansion for the spared whisker. (**B**) IOS area of the untrimmed/principal whisker (PW; β whisker) in baseline session in *Syngap1^+/+^* (n=9 mice) and *Syngap1^+/+^* mice (n=10 mice). (**C**) Change in IOS of the spared area induced by whisker trimming in *Syngap1^+/+^* (one Sample t test t8=2.94, p= 0.0188; n= 9 mice) and *Syngap1^+/−^* mice (one Sample t test t_9_=2.40, p=0.0398; n= 10 mice). (**D**) Same as in (A) but for the trimmed/deprived PW showing no significant changes in IOS area after whisker trimming. (**E**) IOS area of deprived PW in baseline session in *Syngap1^+/+^* and *Syngap1^+/−^* mice. (**F**) Change in IOS of the deprived area induced by whisker trimming in *Syngap1^+/+^* and *Syngap1^+/−^* mice. Bars represent mean ± SEM. Medians are represented by red or black dashed lines. Circles are animal means. *p ≤ 0.05.

**Figure S3.**
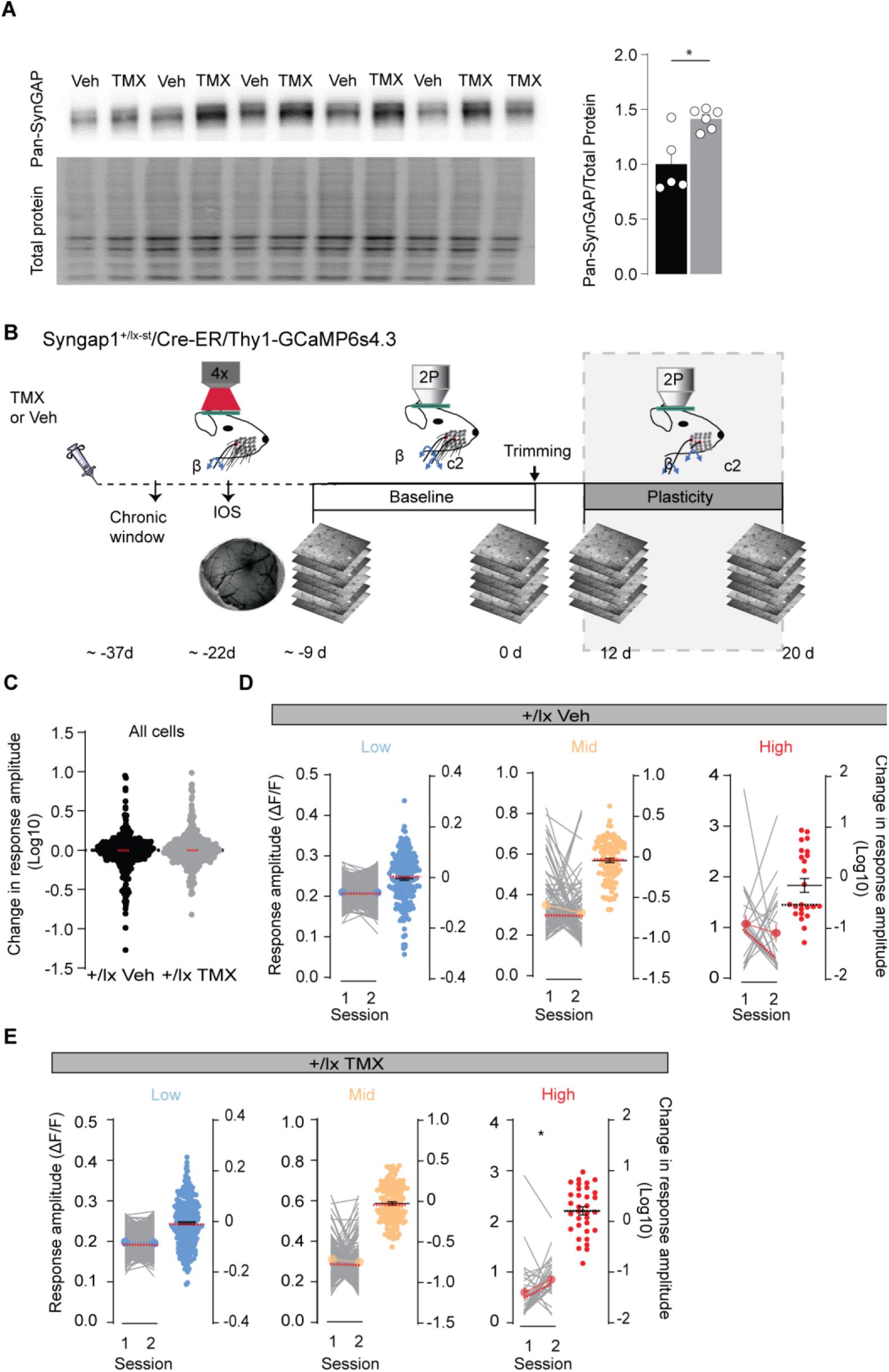
Stable wS1 L2/3 neuron population responsiveness during baseline recording sessions in adult SynGAP re-expression mice. (**A**) Western blot images and graph confirming SynGAP protein re-expression after tamoxifen (TMX) injection to *Syngap1*+/lx mice (n=5 vehicle-injected mice and n=6 TMX-injected mice; Mann Whitney test U=3, p= 0.0303). (**B**) Experimental timeline/design. *Syngap1^+/lx-st^*/Cre-ER/Thy1-GCaMP6s mice were IP injected with tamoxifen (TMX) or vehicle (Veh) and thirty days later implanted with a chronic cranial window and recovered for at least two weeks before Intrinsic Optical Signal (IOS) imaging. IOS was used to identify the cortical locations activated by the deflection of the principal whisker (PW). Baseline imaging sessions (grey shaded region) were carried out ~-9 and 0 days before whisker trimming. (**C**) Change in untrimmed/principal whisker (PW) response amplitude over two baseline sessions for all cells in the spared cortical area in vehicle-(Veh) injected mice (n= 345 neurons from 5 mice) and Tamoxifen-(TMX) injected mice (n= 459 neurons from 6 mice). (D-E) ΔF/F (± SEM) for each cell class over two baseline sessions in (**D**) Veh-injected mice (n=219 low-active cells, n=102 mid-active cells, n=24 high-active cells from 5 mice) and (**E**) TMX-injected mice (n=291 low-active cells, n=135 mid-active cells, n=33 high-active cells from 6 mice, Wilcoxon matched-pairs signed rank test, p=0.0293). Bars represent mean ± SEM Circles are (A) animal mean value, (C) individual cell values and (D and E) cell class mean values. Medians are represented by red or black dashed lines. Lines represent individual cell values. *p ≤ 0.05.

**Figure S4.**
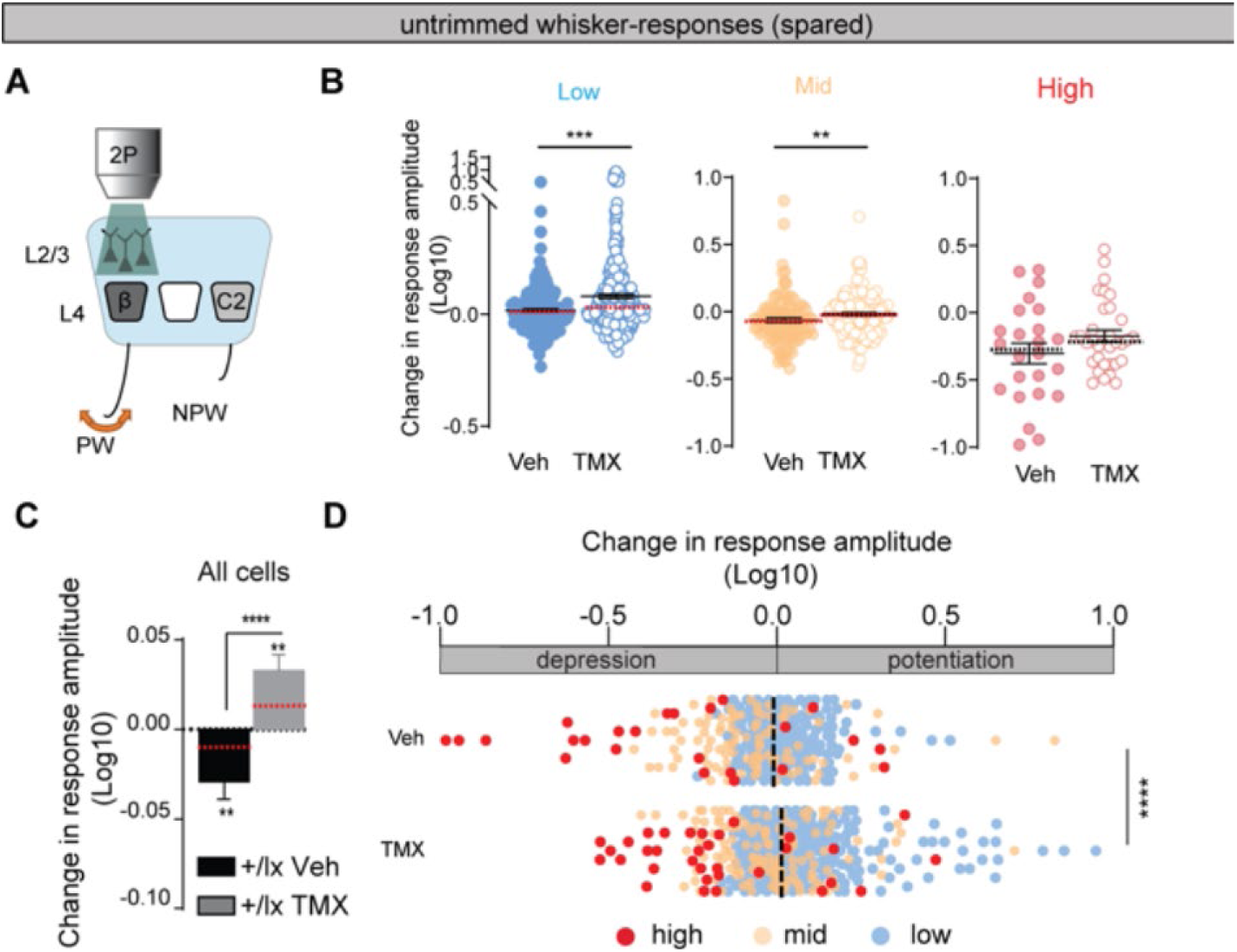
Adult re-expression of *Syngap1* reactivates experience-dependent reorganization of spared synaptic inputs in spared wS1. (**A**) Representation of the experimental design consisting in deflecting the untrimmed/principal whisker (PW) and imaging in the spared area of wS1 from *Syngap1^+/lx-st^*/Cre-ER/Thy1-GCaMP6s mice. (**B**) Change in untrimmed/PW-response amplitude after single whisker experience (SWE) for each cell class in the spared wS1 of Veh-treated mice (n= 219 low-active cells, 102 mid-active cells, 24 high-active cells from 5 mice) and TMX-treated mice (n = 291 low-active cells, 136 mid-active cells, 33 high-active cells from 6 mice; Mann Whitney test U= 25478, p=0.0001; Mann Whitney test U=5523, p=0.0072). (**C**) Change in untrimmed/PW-response amplitude after SWE for all cells in the spared wS1 in Veh-treated mice (n= 345 neurons from 5 mice; Wilcoxon Signed Tank Test, p=0.004) and TMX-treated mice (n= 460 neurons from 6 mice; Wilcoxon Signed Tank Test, p=0.003; Mann Whitney test U=65930, p< 0.0001). (**D**) Bar graphs showing potentiation and depression of all three populations after SWE in Veh-treated (n= 219 low-active cells, 102 mid-active cells, 24 high-active cells from 5 mice) and TMX-treated mice (n = 291 low-active cells, 136 mid-active cells, 33 high-active cells from 6 mice; Mann Whitney test U=65930, p< 0.0001). Bars represent mean ± SEM. Medians are represented by red or black dashed lines. Circles are individual cell values. **p ≤ 0.01, ***p ≤ 0.001 and ****p ≤ 0.0001

## Data and statistics related to main figures

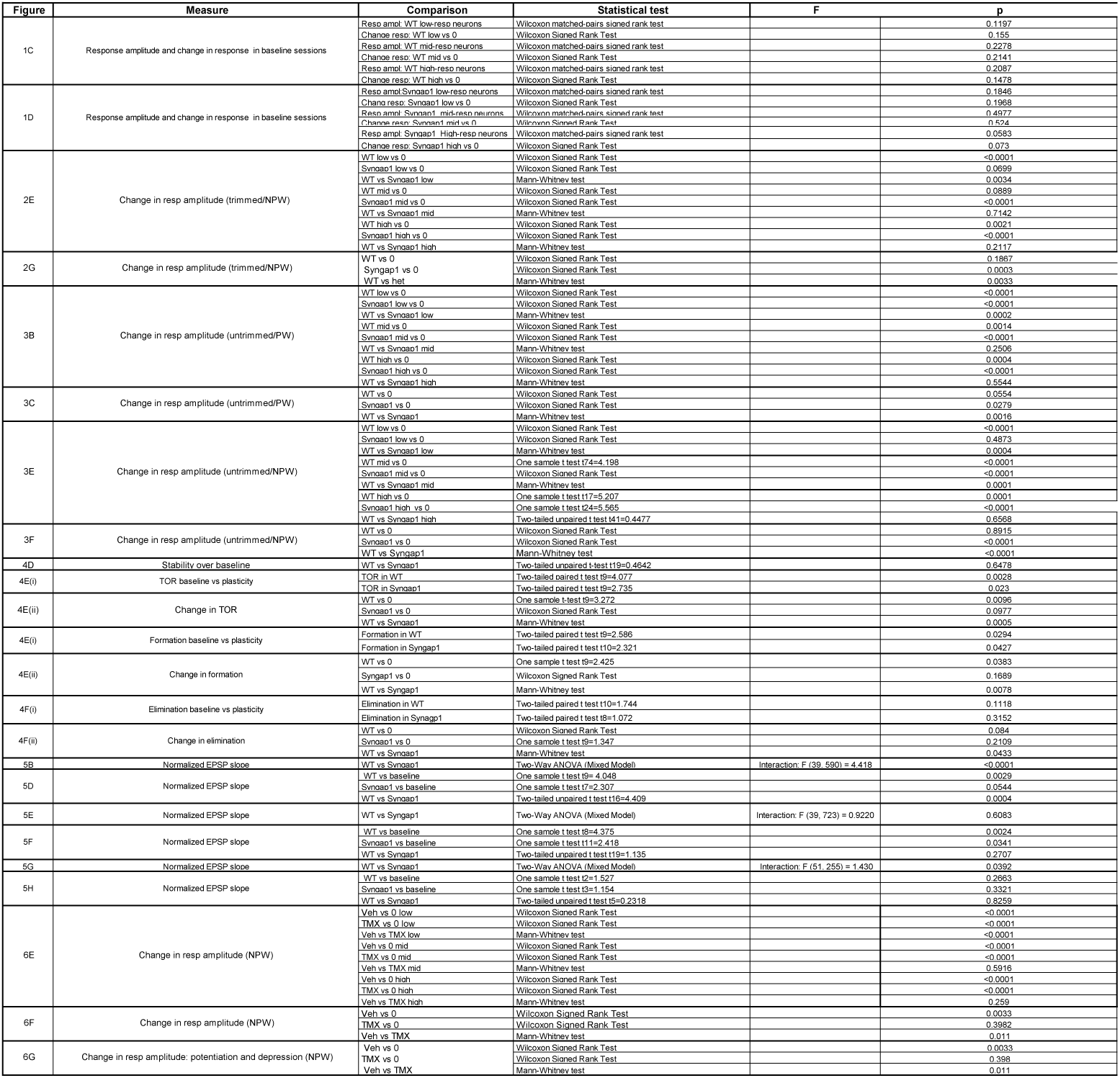

## Data and statistics related to supplementary figures

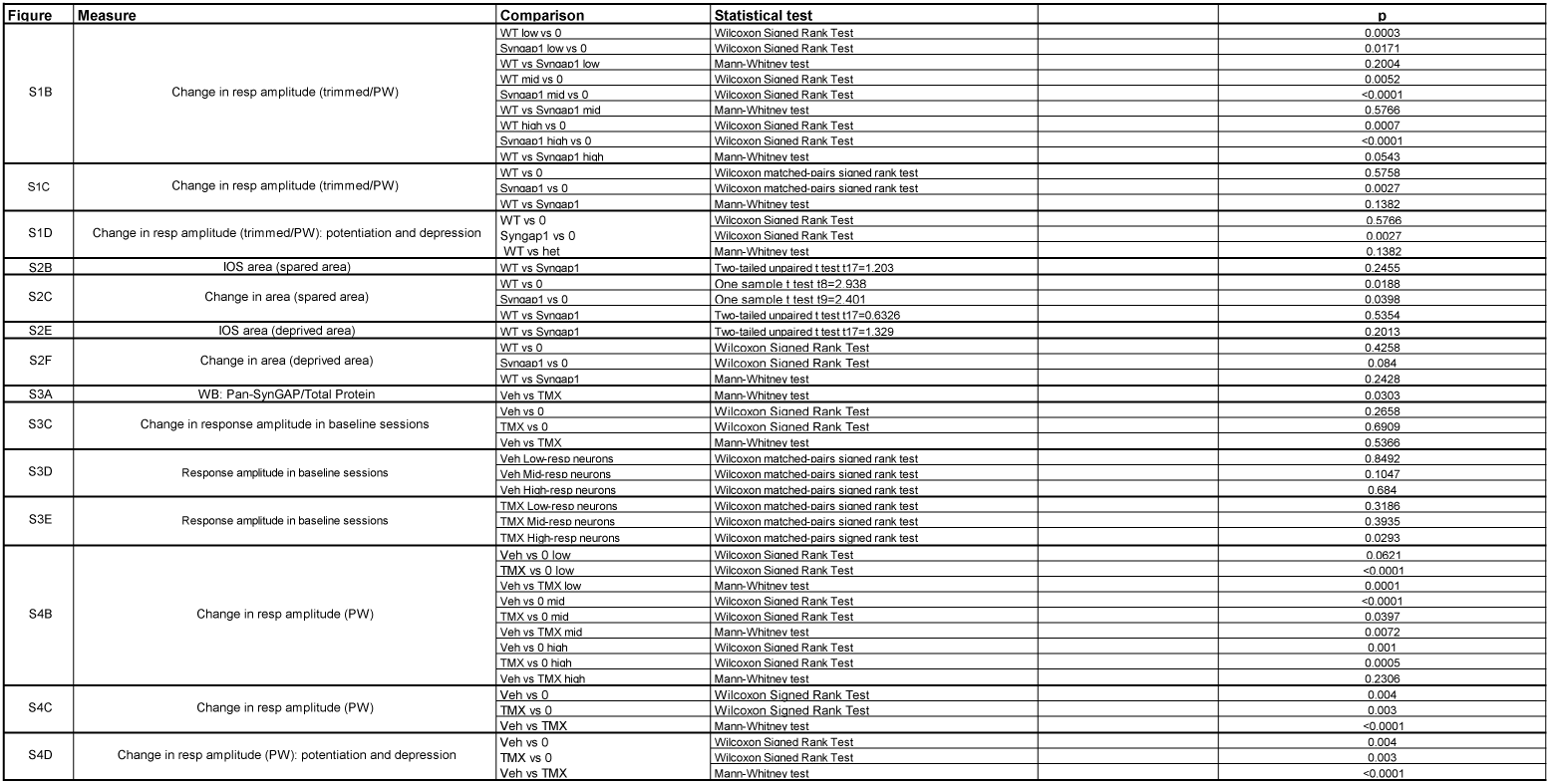

